# Genetic architecture of the *S*-locus supergene revealed in a tetraploid distylous species

**DOI:** 10.1101/2025.04.01.646584

**Authors:** Zhonglai Luo, Spencer C.H. Barrett, Tieyao Tu, Zhongtao Zhao, Shanshan Jia, Shiran Gu, Tingting Duan, Yu Zhang, Bingqiang Xu, Lei Gu, Xiaofang Deng, Libo Jiang, Miaomiao Shi, Dianxiang Zhang

## Abstract

**Summary:**

- Heterostyly is a polymorphic floral adaptation controlled by supergenes and maintained in populations by disassortative mating. The molecular basis of distyly has been investigated in several unrelated families. Previous studies have been limited to diploid species and have not included Rubiaceae, which has the highest number of heterostylous species.
- Here, we investigate the genetic architecture of tetraploid *Schizomussaenda henryi* (Rubiaceae) to identify candidate style-length genes, evidence for hemizygosity and the potential role of gene duplication in the evolution of the *S*-locus. Chromosome-level genome assemblies and transcriptomic approaches identified the *S*-locus region comprised of four hemizygous genes.
- The candidate gene *SchzAUX22* may regulate style length and filament growth by acting in the auxin signaling pathway. Comparative studies of the two sub-genomes of *S. henryi* indicated strong similarities and no significant sub-genome dominance, as expected from autotetraploidy. Evidence from haplotype-resolved assemblies of a short-styled plant indicated that it possessed the *ssSS* genotype, consistent with autotetraploidy, which was estimated to have originated ∼3.6 Mya. Analysis of *Ks* values between *S*-locus genes and their paralogs indicated that the *S*-locus likely originated through stepwise duplication.
- Our study provides comprehensive genomic analysis of distyly in a polyploid and demonstrates that despite whole genome doubling the *S*-locus supergene remains intact and functional as an outbreeding mechanism. Our results contrast with many other heterostylous lineages where polyploidy is associated with breakdown of the polymorphism and transitions to self-fertilization.

## Introduction

Heterostyly is a sexual polymorphism in which populations of animal-pollinated plants are subdivided into two (distyly) or three (tristyly) mating groups characterized by the reciprocal placement of stigmas and anthers between flowers of the floral morphs (Darwin, 1877; Ganders, 1979; Barrett, 1992). The polymorphism is usually associated with heteromorphic self-incompatibility (Barrett and Cruzan, 1994) and ancillary polymorphisms of pollen and stigmas (Dulberger, 1992). Heterostyly has been the subject of sustained interest since Darwin’s seminal work on the topic (Darwin, 1877; Weller, 2009; Cohen, 2010; Gilmartin, 2015) and is reliably reported from 28 angiosperm families and has had numerous origins (Lloyd and Webb, 1992a; Naiki, 2012; Barrett et al., 2000). Today the polymorphism is a paradigmatic model system for studies on the ecology, genetics, and evolution of plant reproductive systems (Scharman and Lenhard, 2024). Experimental studies have confirmed Darwin’s ‘cross-promotion’ hypothesis that heterostyly functions to promote animal-mediated cross-pollination among floral morphs resulting in disassortative mating and maintenance of the floral polymorphism by negative frequency-dependent selection (Lloyd and Webb, 1992b; Barrett, 2019; Simon-Pórcar et al., 2024). Until recently, little was known about the underlying molecular genetic basis of these complex floral polymorphisms, but genomic approaches can now provide novel insights on this problem (reviewed in Kappel et al., 2017).

Studies of the inheritance of distyly in unrelated species have shown that in most cases the polymorphism is under the control of a single diallelic Mendelian locus (S), with dominance of the *S*-allele (short-styled morph – *Ss*; long-styled morph – *ss*; hereafter S- and L-morphs) (Lewis and Jones, 1992). To explain the multiple intermorph differences in structure and physiology in *Primula*, Darlington and Mather (1949) proposed that distyly was associated with a chromosomal region in which a cluster of tightly linked genes protected from recombination governed the polymorphisms. This linkage group is inherited as a single unit and is now referred to as a ‘supergene’ (Thompson and Jiggins, 2014; Schwander et al., 2014; Brennan, 2017). The supergene hypothesis has now been confirmed in several unrelated distylous species (reviewed in Kappel et al., 2017; Barrett, 2019).

Genomic studies on distylous genera in six families, summarized in Table S1, have revealed that the genes comprising the distyly supergene are hemizygous and present only in the S-morph (Li et al., 2016; Huu et al., 2016; Shore et al., 2019; Huu et al., 2020; Gutiérrez-Valencia et al., 2022; Potente et al., 2022; Zhao et al., 2023; Yang et al., 2023; Fawcett et al., 2023). Candidate genes have been proposed in these families but have been studied in most detail in Primulaceae. In Primula 4-5 genes have been identified in the distyly linkage group and gene function has been demonstrated for *CYP^T^*, which determines short styles and incompatibility, and *GLO^T^* which controls anther height (Huu et al., 2016, 2020). Although there is evidence indicating molecular convergence among unrelated distylous species in gene function associated with the inactivation of brassinosteroid (BR) metabolism influencing style length (Li et al., 2016; Shore et al., 2019; Gutiérrez-Valencia et al. 2022; Zhao et al. 2023), other candidate genes have also been identified (e.g. Yasui et al., 2012, Liu et al., 2024). It seems likely, that when a broader sampling of taxa is investigated, a multiplicity of molecular mechanisms are likely to be revealed because of the polyphyletic origins of heterostyly,.

The family with by far the largest number of heterostylous genera and species is Rubiaceae (Bir Bahadur, 1968; Ganders, 1979), which also exhibits a range of derived conditions associated with the evolutionary breakdown of distyly, including homostyly, monomorphic long-styled or short-styled populations, and functional dioecy (Naiki and Kato, 1999; Li et al., 2010; Chen et al., 2013; Duan et al., 2018). This reproductive variation provides opportunities to explore the genetic control of distyly and transitions between sexual systems. However, in Rubiaceae the molecular basis of distyly remains largely unknown, partly due to the lack of whole-genome data for distylous species. Recently, a *S*-locus region in the distylous diploid *Mussaenda lancipetala* has been identified including potential candidate genes regulating style length (Yuan S. et al, unpublished data).

Whole genome duplication (WGD) or polyploidization is the process by which an organism’s entire genetic information is duplicated, once or multiple times. WGD represents a widespread phenomenon in flowering plants influencing evolutionary processes and diverse aspects of reproductive biology (Otto and Whitton, 2000; Soltis and Soltis, 2016; Leslie and Mander, 2024). The extent to which polyploidy might affect *S*-locus structure and function has not been investigated in a heterostylous species but could have important genomic consequences. For example, genomes in allopolyploids, because of their hybrid origin, often exhibit more pronounced genomic divergence with one sub-genome exhibiting greater gene retention, higher gene expression and more conservative evolution, often termed “sub-genome dominance” (Buggs et al., 2012; Yoo et al., 2013 Wang et al., 2022). In contrast, autopolyploids because they result from genome duplication within a species are less likely to evolve sub-genome dominance.

Among heterostylous plants, polyploidization is also commonly associated with the breakdown of distyly to homostyly and the evolution of predominant self-fertilization (Kelso, 1992; Richards 2003; Naiki and Nagamasu, 2004; Guggisberg et al., 2006; Naiki, 2012; Mora-Carrera et al., 2025). However, this association is by no means universal and there are examples of polyploid heterostylous species and also diploid homostyles (Barrett, 1988; Schoen et al., 1997; Shore et al., 2006; Ferraro et al., 2020). An intriguing pattern is evident among some *Turnera* species in which the type of polyploidy maybe be important in determining whether distyly is maintained or not, with autotetraploids that maintain distyly and allopolyploids associated with breakdown to homostyly (Barrett and Shore, 1987; but see Shore et al., 2006). How general the association is between autopolyploidy and the maintenance of heterostyly is unclear. Investigations on the genomic underpinnings of distyly have been conducted exclusively in diploid species and therefore one of the main goals of our study was to investigate the genetic architecture of distyly in a polyploid species, determine the type of polyploidy that occurs, and compare our findings with earlier studies on diploid distylous species.

Here, we present high-quality genome assemblies and transcriptome sequencing of distylous tetraploid, *Schizomussaenda henryi* (Rubiaceae). This species exhibits a typical distylous syndrome characterized by reciprocity between stigma and anther heights between the floral morphs (Jia, 2024) and a well-developed self- and intra-morph physiological incompatibility system (Deng, 2007). Our investigation specifically addressed the following questions: (1) Using genomic and transcriptome data can we locate the *S*-locus supergene region of *S. henryi*, identify potential candidate genes governing sex-organ dimorphism, and determine if *S*-locus genes are hemizygous? These objectives were of importance in assessing the extent to which there was evidence of molecular convergence between Rubiaceae and other heterostylous families; (2) Is *S. henryi* an autopolyploid or an allopolyploid and is there evidence of sub-genome dominance? We predicted that if the species is allopolyploid some degree of sub-genome dominance should be evident but if *S. henryi* is autotetraploid we expected no significant sub-genome dominance. (3) Has WGD played a role in the evolutionary history of *S*-locus genes and is there evidence that stepwise gene duplication may have promoted the evolutionary assembly of the supergene, as reported in several other distylous species (e.g. Huu et al., 2020; Potente et al., 2022).

## Results

### Karyotype analysis and genome size estimation

The L- and S-morphs *S. henryi* each had 44 chromosomes (Fig. S1B, C) consistent with tetraploidy. This chromosome number is identical to tetraploid *Coffea arabica* (2n=4x=44, Scalabrin et al. 2024) and double the number of chromosomes in diploid *Coffea canephora* and *Mussaenda shikokiana* (2n=2x=22, Denoeud et al., 2014; Chen, 2013). The genome size of the S-morph of *S. henryi* was ∼650 Mb estimated by flow cytometry, a value slightly smaller compared to the *k*-mer based analysis (∼700 Mb, *k*=21, p=4).

### Genome sequencing, assembly, and annotation

The sizes of assembled genomes for the L- and S-morph were 743.1 and 720.4 Mb, with scaffold N50 of 10.35 and 12.94 Mb, respectively. The BUSCO completeness values (C) exceeded 98.0 % for both morphs (98.6 % and 98.8 % for the L- and S-morph respectively), with only 0.9% missing BUSCO genes, indicating a high completeness of the assemblies (Table S2). With the aid of Hi-C technology, we anchored contigs to 22 chromosomes (pseudomolecules, Chr01-Chr22) (Fig. 1 and S2), and the chromosome-level genome was 705 and 715Mb for the L- and S-morphs, respectively. The principal features of the assembled genome are illustrated in Circos plots (Fig. 1).

**Figure 1.**
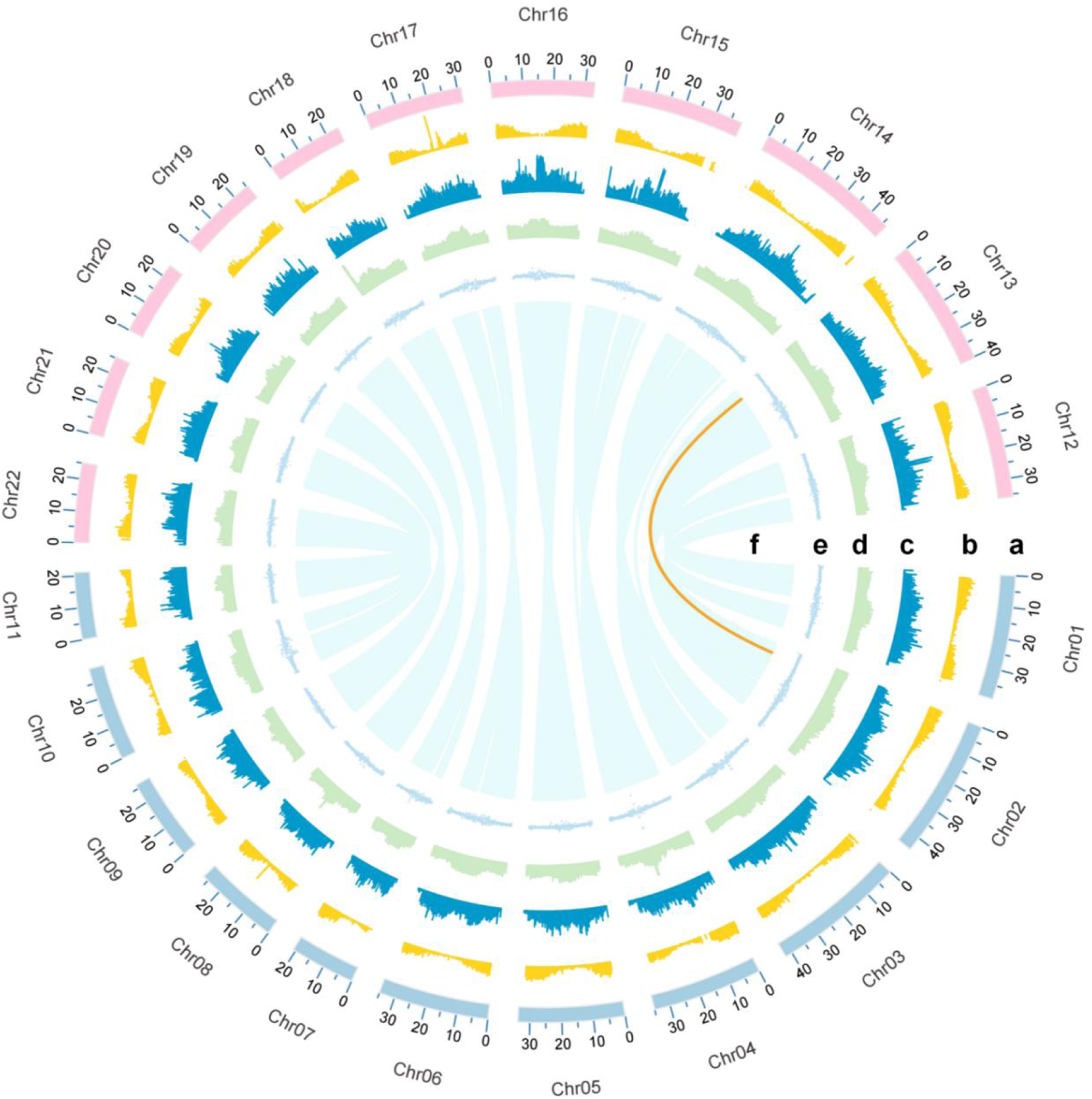
Circos map illustrating basic information on the *Schizomussaenda henryi* genome (S-morph). a, chromosome information; b, gene density; c, TE density; d. GC content; e, GC skew; f, chromosomal collinearity within genome. Sub-genome A (Chr01 - Chr11) and sub-genome B (Chr12-Chr22) are highlighted in light-blue and pink, respectively. The collinearity between the genes at the upstream and downstream boundaries of the S-locus on Chr13 and their homologous genes on Chr02 is indicated by orange lines.

We used chromosome-level sequences for annotation of repetitive elements and genes. RepeatMasker annotation revealed that 54.7% and 53.17 % of the L- and S-morph genomes, respectively, consisted of repeats, with LTR accounting for the highest proportion (ca. 40 %). These sequences were further categorized into various subtypes (Table 1). In all, 51,955 and 52,600 protein-coding genes were predicted for the L- and S-morph genomes, respectively. BUSCO evaluation based on the “protein” model revealed completeness scores (C) of 96.5% and 96.6% for L- and S-morph gene predictions, respectively. The average length of coding genes was 3,057 bp, with an average of 4.72 exons per gene.

**Table 1.** Repeat sequence annotations for the genome of the long- and short-styled morphs of *Schizomussaenda henryi*.

| Category | No. of elements |  | Length (bp) |  | Percentage (%) |  |
| --- | --- | --- | --- | --- | --- | --- |
|  | L-morph | S-morph | L-morph | S-morph | L-morph | S-morph |
| DNA transposons | 71,270 | 68,956 | 19,258,895bp | 18,424,182 bp | 2.59% | 2.62 % |
| LINEs | 13,809 | 13,342 | 9,497,875bp | 9,028,995 bp | 1.28% | 1.28 % |
| SINEs | 1,229 | 1,188 | 186,406bp | 179,891 bp | 0.03% | 0.03 % |
| LTR elements | 226,343 | 216,004 | 298,242,833bp | 275,641,539 bp | 40.14% | 39.14 % |
| Rolling-circles | 4,386 | 4,333 | 3,275,765bp | 3,082,096 bp | 0.44% | 0.44 % |
| Unclassified | 256,511 | 247,609 | 70,599,679bp | 66,899,542 bp | 9.50% | 9.50 % |
| Small RNA | 3,091 | 2,914 | 3,446,452bp | 1,123,455 bp | 0.46% | 0.16 % |
| Simple repeats | 760 | 760 | 29,935bp | 29,978 bp | 0.00% | 0.00 % |
| Low complexity | 0 | 0 | 0bp | 0 bp | 0.00% | 0.00 % |
| Total | 577,399 | 555,106 | 404,537,840bp | 374,409,678bp | 54.47% | 53.17% |

We performed gene annotation using the eggNOG-mapper (v.2) pipeline. In the S-morph genome, 45,327 (> 86 %) protein-coding genes were successfully annotated in at least one database (Table S3). A total of 4,293 GO terms in three ontology categories (Biological Process, BP; Molecular Function, MF; Cellular Components, CC) were assigned to 34,500 genes. The top three highly represented GO terms were: “integral component of membrane” (GO:0016021), “ATP binding” (GO:0005524), and “nucleus” (GO:0005634) and overall, 17,573 genes were matched to at least one KEGG pathway (Table S3).

### Sub-genome reconstruction and analyses

Basing on sub-genome-specific *k*-mers, SubPhaser failed to divide the genome into two groups with equal chromosome numbers (Supplementary Data 1). With the aid of phylogenetic methods (see Methods), we assigned the 11 chromosomes, LG12, LG13, LG03, LG04, LG16, LG17, LG07, LG19, LG20, LG21, LG11, which had closer genetic distances to *M. pubescens*, to sub-genome B (named as Chr01-Chr11), and another 11 chromosomes (LG01, LG02, LG14, LG15, LG05, LG06, LG18, LG08, LG09, LG10, LG22) to sub-genome A (named as Chr12-Chr22) (Fig. S3). Syntenic regions between the sub-genomes of S-morph *S. henryi* were identified by JCVI and are illustrated in Fig. 2. The collinearity between the two sub-genomes indicates a high degree of similarity between them.

**Figure 2.**
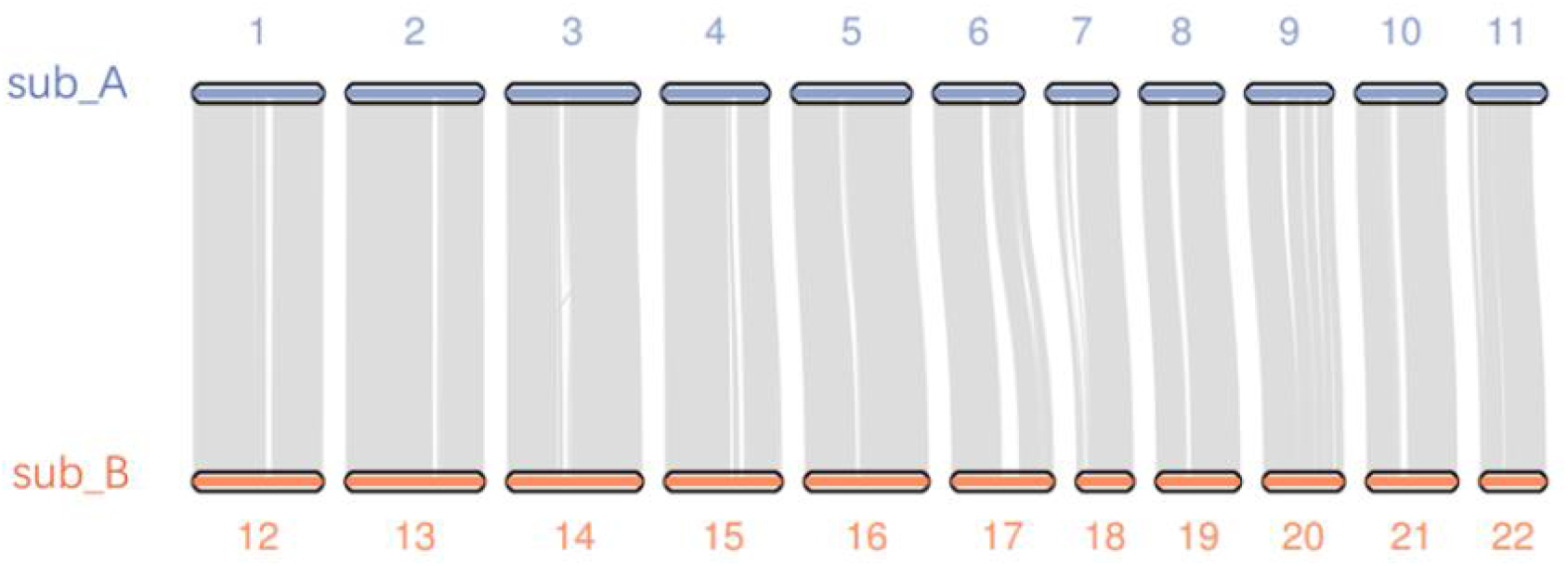
Syntenic analysis of the two sub-genomes of the short-styled morph of *Schizomussaenda henryi.* sub_A, sub-genome A; sub_B, sub-genome B.

For the S-morph, Chr02 had the highest number of genes (3,259 genes, 15,268 exons), whereas Chr07 had the fewest genes (1,310 genes, 6,391 exons). The density of gene distribution on each chromosome is illustrated in Fig. S4. The average gene density was highest on Chr16 (86.64 genes/Mb), followed by Chr05 (83.54 genes/Mb) and Chr12 (83.11 genes/Mb). The chromosome with the lowest gene density was Chr06 (60.92 genes/Mb).

RNA-seq data derived from styles and filaments of two development stages indicated no significant difference in the expression of homoeologous gene pairs between sub-genomes A and B (all the *p*-values > 0.1 in Mann-Whitney Rank Sum Test). We also compared the number of homoeologous gene pairs with biased expression (*p* < 0.01, and over two-fold changes) towards either sub-genome A or B and found no significant difference across different tissues and development stages (*p* = 0.485 in Mann-Whitney Rank Sum Test, Fig. S5).

To evaluate selection on the two sub-genomes, we calculated *Ka* and *Ks* values for homoeologous gene pairs between *C. canephora* and the two sub-genomes of *S. henryi*. We found that every homoeologous chromosome pair showed similar *Ka/Ks* values (∼0.226-0.237, all *p*-values > 0.1 in Mann-Whitney Rank Sum Test; Table S4). There were 96 (4.1%) and 79 (3.5%) BUSCO genes lost in sub-genomes A and B, respectively, 23 of which were identical. Sub-genomes A and B had 73 and 56 sub-genome-specific missing BUSCOs, respectively; Supplementary Data 2). Significantly, the BUSCOs lost from sub-genome A were retained in sub-genome B, and vice versa, which may explain why only 0.9% of BUSCOs were missing from the total genome.

### Whole-genome duplication

The distribution of *Ks* for paralogous genes in the *S*. *henryi* genome showed two prominent peaks at ca. 1.84 and 0.056, respectively, indicating that the species experienced at least two whole-genome duplication events. The ancient WGT (peak I in Fig. 3, *Ks*=1.84) was a conserved whole-genome triplication event that was shared with diploid *C. canephora, M. pubescens and T. quinquecostatus*, which was estimated to have occurred at ca. 118.3 -124.7 Mya. The second and more recent peak (peak II) in *S*. *henryi* at *Ks*=0.05596, probably associated with tetraploidization, occurred at ∼3.6 Mya and did not occur in the three diploid species and was apparently a species-specific duplication event. Significantly, the divergence peak for sub-genomes A and B of *S*. *henryi* (peak with red dashed line) had very similar *Ks* value (0.05606) with peak II (Fig. 3).

**Figure 3.**
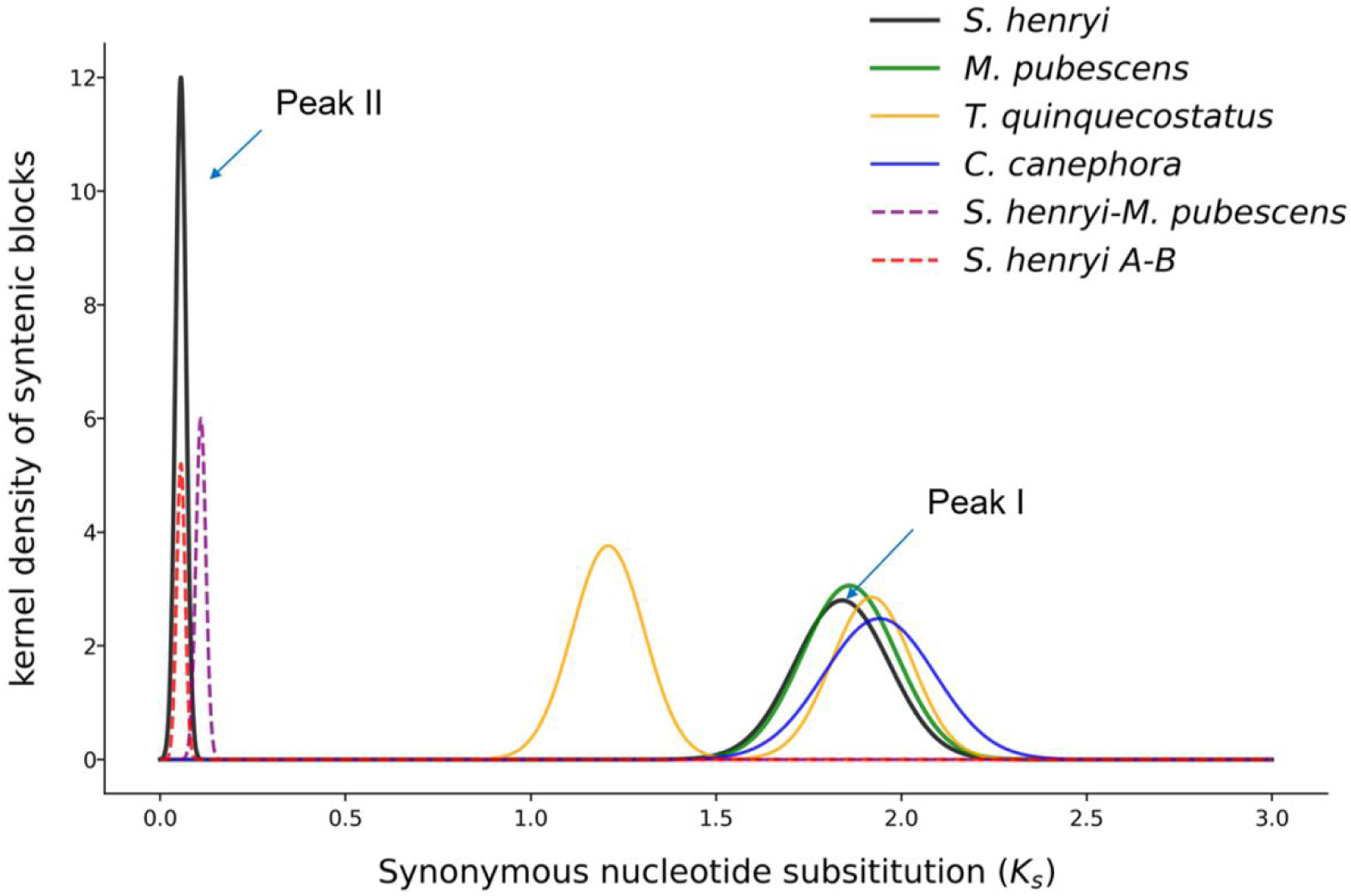
*Ks* distributions of paralogous genes in *Schizomussaenda henryi* (S-morph), *Mussaenda pubescens*, *Coffea canephora* (Rubiaceae) and *Thymus quinquecostatus* (Lamiaceae).

### Identifying the *S*-locus region and detecting hemizygosity

DEGs expressed in the S-morph styles but barely expressed in the L-morph (FPKM<1.0) were selected as possible candidate genes for distyly and their biological functions examined according to annotations. *S13.827*, a homolog of the auxin-induced protein AUX22 gene, plays a role as a repressor of early auxin response genes (Tiwari et al., 2001; Ulmasov et al., 1997). *S13.827* (*SchzAUX22*) was highly expressed in S-morph styles at both early (FPKM value = 256.6) and late (FPKM value = 471.4) stages, whereas no expression was detected in the L-morph (Fig. 4A-D). Another gene, *S13.824*, located upstream of *S13.827*, is a gibberellin 3 beta-dioxygenase (*GA3ox*). Its homolog *GeGA3ox* has been suggested in the control of anther position in distylous *Gelsemium elegans* through regulating the synthesis of bioactive gibberellins (Zhao et al., 2023). In S-morph styles of *S*. *henryi, S13.824* (*SchzGA3ox*) exhibited low expression in the early developmental stage, and no expression in the later stage. This result suggests that the gene might be expressed transiently in the early period of style development. No expression was observed in L-morph styles (Fig. 4E, F).

**Figure 4.**
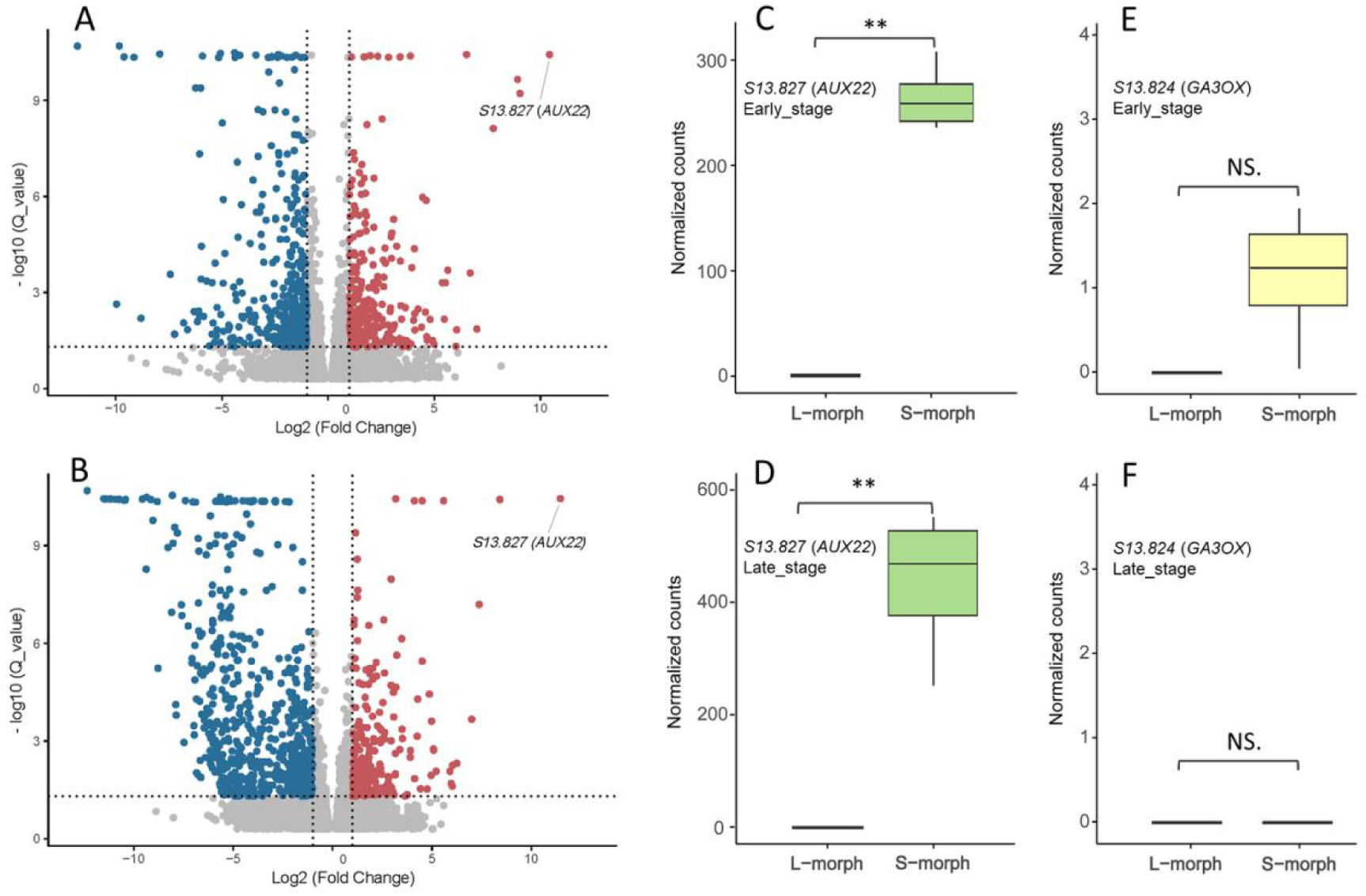
Differential expression of *S*-locus genes *S13.824* and *S13.827* in the styles of *Schizomussaenda henryi*. Normalized counts of both genes between L- and S-morph styles are compared. ** *p* < 0.01 in Wilcoxon rank-sum test. NS. Difference not significant. A, C, E, early stage; B, D, F, late stage.

We compared the genome sequences containing *S13.824* and *S13.827* between the S- and L-morph with the aid of IGV. We identified a ∼75 kb region (from ca. 5,562 kb to 5,637 kb) on Chr13 of the S-morph genome, which was absent from Chr13 of the L-morph genome. Sequence coverage for the region was about half of that for the flanking areas indicating this region was hemizygous (Fig. S6).

After manual curation, we found that the region contained four annotated genes, *S13.822, S13.824, S13.826,* and *S13.827*, none of which were evident in the L-morph genome (Fig. 5). The functions of *S13.827* and *S13.824* and the hemizygous structure of the S-locus region are consistent with the hypothesis that this region contains candidate *S*-locus genes governing distyly in *S*. *henryi*. Functional annotation indicated that *S13.822* is a *pectin acetylesterase 5* (*PAE 5*)*-like* gene, whereas *S13.826* is *DEAD-box ATP-dependent RNA helicase 37* (*RH37*)*-like* gene. However, these two genes (*S13.822* and *S13.826)* did not express in either the style or filament-floral tube tissues of the S-morph (FPKM value = 0). To further examine the presence-absence variation at the *S*-locus between floral morphs, we mapped Illumina sequences generated from wild populations to the genome assemblies. Genome resequencing data mapping confirmed that the *S*-locus was S-morph specific, with the four *S*-locus genes all absent from L-morph individuals (Fig. S7).

**Figure 5.**
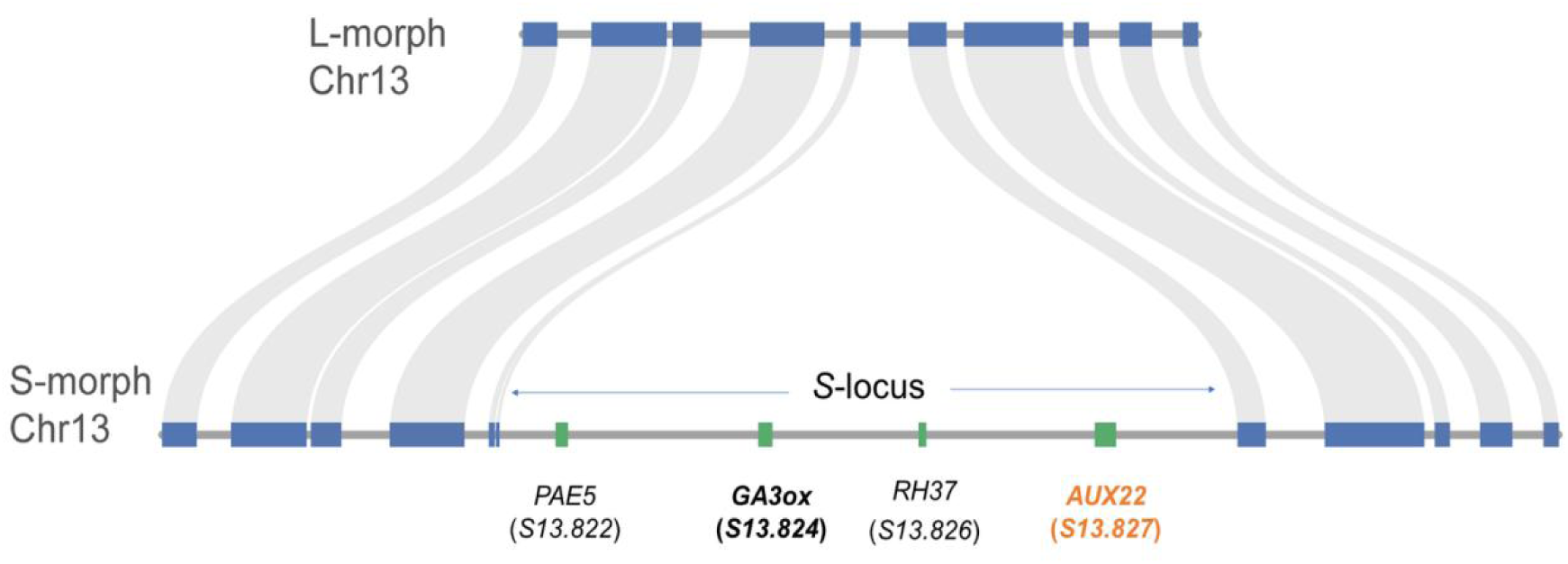
Model of the genetic architecture of *S*-locus genes in *Schizomussaenda henryi*.

*S*-locus genes reported from distylous *Primula, Gelsemium, Turnera, Linum*, and *Fagopyrum* were selected for BLAST analysis against the L- and S-morph genomes of *S. henryi* and we compared the expression of homologous genes between the morphs. There were no significant differences in expression of these *S*-locus homologs between L- and S-morph styles or filaments in *S. henryi*. This finding indicating their presence in both floral morphs strongly suggests they are not involved in the control of floral dimorphism in *S. henryi*.

### Origin and evolution of *S*-locus genes

To examine if the *S*-locus genes originated through segmental duplication or stepwise duplication, we searched for paralogs of *S*-locus genes in the S-morph genome of *S*. *henryi*. *S13.827* and *S10.1705* on Chr10 were identified as the closest paralogs. Moreover, paralogs for the *S*-locus genes *S13.822,* and *S13.824* were all located on Chr10 with a scattered distribution (*S10.709,* and *S10.1697*, respectively) (Fig. 6). Significantly, the closest paralog for *S13.826* was found on Chr14 (*S14.2172*). In contrast, the paralogs of genes adjacent to the *S*-locus (i.e., upstream from *S13.822*, and downstream from *S13.827*) were arranged in order on Chr02, which is homologous to Chr13. This arrangement supports the hypothesis that the *S*-locus genes were originally inserted into Chr13 from other chromosomes.

**Figure 6.**
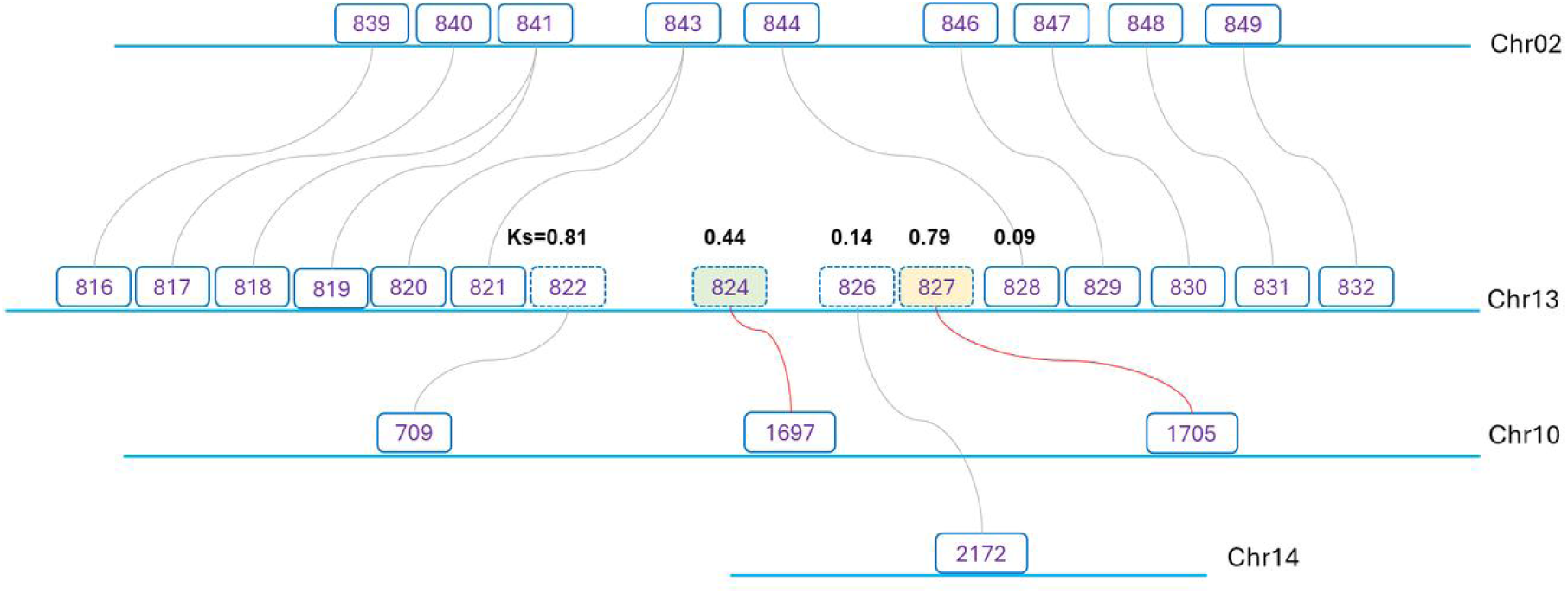
Schematic plot showing homologous genes of the *S*-locus in the genome of the short-styled morph of *Schizomussaenda henryi* with respective *Ks* values.

We calculated *Ks* values between S-locus genes and their paralogs because if the *S*-locus supergene originated by stepwise duplications the genes should exhibit different ages (see Potente et al., 2022; Yang et al., 2023). Our results showed that genes at the left and right ends of the *S*-locus had higher *Ks* values of 0.81 for *S13.822*/*S10.709*, and 0.79 for *S13.827*/*S10.1705*. In contrast, *S13.826*/*S14.2172* had the lowest *Ks* of 0.14. The *Ks* values and their flanking genes and paralogs are illustrated in Fig. 6 which shows that they descend “centripetally” consistent with stepwise duplications, with *S13.822* first, *S13.827* second, followed by *S13.824*, and *S13.826* last.

We computed Evolutionary Distances (EDs) of *S*-locus flanking regions (including both the genes and non-coding sequences) between L- and S-morph genomes. We observed prominent K2P peaks at approximately 200-1000 bp upstream and 200-400 bp downstream of the hemizygous region indicating genetic differentiation increased in regions near the *S*-locus (Fig. S8).

TEs (transposable elements) are often enriched in *S*-locus regions of distylous species, (e.g. Shore et al., 2019; Yang et al., 2023; Zhao et al., 2023). The density of TEs (based on 5 kb windows) was significantly higher across the *S*-locus, from approx. 5570 kb to 5630 kb, compared to flanking regions approx. 60 kb upstream (*p* < 0.01 in Mann-Whitney Rank Sum Test) and 60 kb downstream (*p* < 0.001 in Mann-Whitney Rank Sum Test, Fig. S9). In addition, the average TE density of the *S*-locus region (1.51 TEs per kb) was ca. 41% higher than that of the rest of Chr13 (1.07 TEs per kb). We also analyzed the types of TEs near hemizygous genes and a MuDR element (‘Mutator Don Robertson’, Robertson’s Mutator) was identified just downstream of *S13.824* (*SchzGA3ox*).

### Genetic structure of *S*-locus and molecular evidence for autopolyploidy

Our comparative analyses of the two sub-genomes (haplotypes) of *S*. *henryi* revealed high similarities between sub-genomes A and B suggesting an autopolyploidy origin. In autotetraploid distylous species, genotypes could be either *SSss* or *Ssss* for the S-morph and *ssss* for L-morph. To examine the genetic structure of *S*-locus, we employed HiFiasm to perform a haplotype-phased assembly for the S-morph genome by setting the parameter --n-hap 4. Then we made BLAST searches using the *S*-locus gene sequences against the four assemblies. The four genes (*S13.822, S13.824, S13.826,* and *S13.827*) had 100 % matches on ctg0046 of hap3 and ctg0045 of hap4 whereas no hits were found in hap1 and hap2. This result is consistent with the hypothesis that the sequenced S-morph individual was of genotype *SSss* (Supplementary Data 3).

## Discussion

We constructed high-quality chromosome-level assemblies of distylous *S. henryi* (Rubiaceae) and characterized its tetraploid genome. Comparisons of the two sub-genomes indicated little evidence of sub-genome dominance, consistent with the autotetraploid status of the species. Evolutionary and WGD analyses indicated that the formation of the *S*-locus likely occurred earlier than the polyploidization event. A hemizygous region in the S-morph genome containing *SchzAUX22* was identified as a possible candidate gene regulating style length. To our knowledge, this is the first investigation at the genomic level of the *S*-locus supergene governing distyly in a polyploid species. We now discuss the genomic features of distyly in *S. henryi* and the evolutionary formation and potential function of *S*-locus genes.

### The tetraploid genome of *S. henryi*

Phylogenetic studies indicate that monotypic *Schizomussaenda* is one of the closest relatives of *Mussaenda* in the tribe *Mussaendeae* (Ixoroideae), and their ancestors diverged approximately 14.7 Mya (Duan et al., 2008). All species of *Mussaenda* that have chromosome records (data from IPCN Chromosome Reports http://legacy.tropicos.org/Project/IPCN and Chen, 2013) are diploid with 22 chromosomes. Our cytological studies demonstrated that *S. henryi* is tetraploid with 44 chromosomes and its genome size was about twice that of the diploid *Mussaenda pubescens* (∼390 Mb, Jia, 2024; Luo et al., unpublished data) or *Coffea humblotiana* (∼422 Mb, Raharimalala et al., 2021).

Sub-genome dominance has been reported in various polyploid plants and animals. For example, in the allotetraploid *Orychophragmus violaceus* (Brassicaceae), higher gene density and expression levels were reported in sub-genome OV1 than OV2 (Zhang et al., 2023). Similar trends have been reported in *Juglans regia* (Juglandaceae) with biased gene expression evident between sub-genomes in various tissues (Li et al., 2023). We compared the expression of homoeologous gene pairs between sub-genomes in style and filament tissues of *S. henryi* and found no significant differences, a result consistent with autotetraploidy.

Different sub-genomes in polyploid species may experience contrasting selective pressures. For example, in the allotetraploid fish *Cyprinus carpio* (Cyprinidae), the mean *Ka*/*Ks* values were significantly lower in the dominant sub-genome B compared to sub-genome A suggesting that genes in sub-genome B have undergone greater purifying selection than their homoeologs in sub-genome A (Xu et al., 2019). We compared the *Ka*/*Ks* values of homoeologous gene pairs between *C. canephora* and the two sub-genomes of *S. henryi* and found no significant difference between each homoeologous chromosome pair (*p*-values > 0.1 for the 11 homoeologous chromosome pairs). These results revealed no obvious sub-genome dominance in *S. henryi* and the two sub-genomes have likely experienced similar selective pressures. The high similarity between sub-genomes in our syntenic analysis (Fig. 2) provides further evidence that *S. henryi* originated from an auto-tetraploidization event from a single diploid ancestral species.

The proportion of missing BUSCO genes from sub-genomes A and B of *S. henryi* was significantly reduced compared with allotetraploid *Acorus calamus* (Ma et al., 2023) and allohexaploid *Carassius gibelio* (Kuhl et al., 2022). Nevertheless, complementarity of the sub-genomes was evident with 17 (ca. 23.3%) fewer missing BUSCOs in sub-genome B compared to A (Supplementary Data 2) suggesting that sub-genome B may exhibit more conservative genome evolution.

### Structure and potential function of *S*-locus genes in hormone control

The molecular mechanism controlling stylar dimorphism was first reported in *Primula* (Li et al., 2016; Huu et al., 2016) and has subsequently been studied in a handful of families within the past decade (e.g. Shore et al., 2019; Zhao et al., 2023; Yang et al., 2023 Liu et al., 2024). In all cases, the *S*-locus is hemizygous and candidate genes have been identified that likely function in regulating style length by inactivating brassinosteroids in *Primula* and *Turnera* (Huu et al. 2016; Matzke et al. 2021 and see Table S1). We identified a ∼75 kb segment on Chr13 of the S-morph of *S. henryi* as the potential S-locus region. Genome annotation and structure analyses revealed that this region was also hemizygous and present only in the S-morph (Fig. 5 and S6). Mapping of genome resequencing data further supported that this region was S-morph specific (Fig. S7).

Four annotated genes occurred at the *S*-locus of *S. henryi*: *S13.822* (*SchzPAE5*), *S13.824* (*SchzGA3ox*), *S13.826* (*SchzRH37*), and *S13.827* (*SchzAUX22*) (Fig. 5). High expression of *SchzAUX22* was only detected in S-morph styles at both early and late stages. Its homolog, auxin-induced protein *AUX22*, was first characterized in soybean (*Glycine max*) (Ainley et al., 1988). AUX/IAA proteins are short-lived transcriptional factors that function as repressors of early auxin response genes. Studies in model plants have demonstrated all the Aux/IAA proteins has repressive effects on the transcription of auxin-responsive reporter genes (Tiwari et al., 2001; Ulmasov et al., 1997). Ectopically overexpression of *OsAUX/IAA* results in dwarfism and decreased gravity response in transgenic rice (Song & Xu, 2013). Down regulation of *AUX/IAA2* in potato (*Solanum tuberosum*) significantly increases plant height (Kloosterman et al., 2006). The high expression of *S13.827* (*SchzAUX22*) in S-morph styles may function to suppress elongation through inhibiting the activity of auxin response factors (ARFs).

Auxin plays a pivotal role in a wide range of plant developmental and physiological processes with auxin response genes playing dual roles in regulating cell division and act in a tissue-dependent manner (e.g. Cao et al., 2019; Béziat et al., 2017). Certain ARFs activate genes that facilitate cell division in root meristem, whereas in older tissues they may repress division signals and reduce organ growth (Truskina et al. 2021; Leyser, 2018; Wang et al., 2005). These findings from other plant systems may help explain the high expression of *S13.827* (*SchzAUX22*) in both S-morph styles and filaments. We propose that this gene may function pleiotropically in opposing ways — inhibiting style elongation while also promoting cell division in filaments (corolla tubes). Functional analysis of *SchzAUX22* would be required to test this hypothesis.

Gibberellin (GA) is another important phytohormone that regulates organ growth and development. GA3ox is a key enzyme involved in GA biosynthesis, converting inactive gibberellin precursors into bioactive gibberellins (Sun et al. 2023). In a gibberellin-deficient tomato mutant, stamen growth was retarded, suggesting GA may promote filament elongation (van den Heuvel et al., 2001). In distylous *Gelsemium elegans*, Zhao et al. (2023) reported *GeGA3ox* as a potential candidate gene that regulates filament growth. Our data revealed that *SchzGA3ox* (*S13.824*) located in the *S*-locus region but exhibited very low expressions in early-stage styles and filaments and was not expressed in the late-stage tissues (Fig. 4E, F). This finding may be because: (1) the gene is transiently expressed in a very early period of development that we did not sample; (2) *SchzGA3ox* in *S. henryi* does not function in regulating filament growth. The second hypothesis may be more plausible if *S13.827* (*SchzAUX22*) controls both style length and filament growth in differed directions.

Besides plant hormone-related genes, genes with unknown or poorly characterized functions have also been identified within the *S*-locus of distylous plants (e.g. Yang et al., 2023; Zhao et al., 2023). In addition to *SchzAUX22* and *SchzGA3ox*, two other genes were located at the *S*-locus of *S. henryi*. *S13.822* encodes a homolog of *pectin acetylesterase 5* (*PAE 5*), which may regulate cell wall architecture and developmental processes by modulating acetate levels in cell walls (de Souza et al., 2014). The second gene, *S13.826*, is annotated as a *DEAD-box ATP-dependent RNA helicase 37* (*RH37*)*-like* gene, which has been implicated in stress tolerance and developmental regulation (Huang et al., 2016; Nawaz et al., 2021). Neither gene exhibited detectable expression in the style or filaments of S-morph flowers, however, suggesting potential functional loss. Further investigations are warranted to elucidate their potential roles within the *S*-locus.

In diploid distylous *Mussaenda lancipetala*, we have identified an *S*-locus region with three hemizygous genes present only in the S-morph, which also contains the *AUX/IAA* gene and *GA3ox* gene (Yuan et al, unpublished data). The ancestral *S*-locus governing the distylous floral syndrome has therefore been maintained after auto-tetraploidization. This finding contrasts with the situation in *Primula* where polyploidization is associated with the breakdown of distyly to homostyly (Kelso, 1992; Richards 2003; Guggisberg et al. 2006). Recent evidence from homostylous, allopolyploid *P. grandis* indicates that the breakdown to homostyly is associated with degeneration of *CYP^T^*, the gene governing style length and female self-incompatibility, likely resulting from relaxed selection and the accumulation of small loss-of-function mutations (Mora-Carrera et al., 2025). Interestingly, the remaining genes in the *S*-locus supergene of *P. grandis* have been largely unaffected by this breakdown event and there was minimal evidence of sub-genome dominance.

### Transposable element accumulation at the S-locus region

TEs can play a significant role in the evolution of genomes by influencing genetic variation, gene regulation and genome architecture (Oliver et al., 2013; Kent et al., 2017; Bourque et al., 2018), and in some cases floral morphology (Kovarik et al., 2008; Fambrini & Pugliesi, 2018). Previous molecular studies of distylous plants have reported the accumulation of transposable elements (TEs) in the *S*-locus region (e.g. *Turnera*, Shore et al., 2019; *Linum*, Gutierrez-Valencia et al., 2022; *Nymphoides*, Yang et al., 2023). Our finding for *S. henryi*, were consistent with these finding since TEs were significantly enriched across the *S*-locus (Fig. S9). In concert with hemizygosity, TEs probably contribute to recombination suppression and hence the maintenance of floral polymorphism.

Additionally, we identified prominent K2P peaks upstream and downstream of the hemizygous region (Fig. S8), providing evidence of elevated molecular divergence between alleles (and see Zhao et al., 2023). Members within supergenes often exhibit high molecular divergence due to strong linkage because of suppressed recombination, a crucial mechanism for the maintenance of heterostyly (Gutierrez-Valencia et al., 2022). In our study, we identified a MuDR element near *SchzGA3ox*. These transposable elements belong to a major type of Class-II transposons first identified in maize (Robertson 1981) and are reported to play central role in gene regulation and genome evolution (Jordan et al., 2003; Kazazian, 2004; Skibbe et al. 2009). Whether this element contributes to suppression of gene expression of certain S-locus genes or is involved in the translocation of genes to the S-locus requires further investigation.

### Evolutionary origins of the *S*-locus and the maintenance of distyly

Two main models have been proposed to explain the evolutionary origins of *S*-locus supergenes (reviewed in Kappel et al., 2017): 1) the segmental duplication model, in which a genomic segment with precursors of *S*-locus genes was duplicated; 2) the stepwise duplication model, in which *S*-locus genes are duplicated and translocated to the same genomic region independently of one another. In *S. henryi,* the closest paralogs of the four *S*-locus genes were distributed on two different chromosomes, with three located on Chr10 and one on Chr14 (Fig. 6). *Ks* values between the *S*-locus genes and their paralogs in the S-morph genome ranged from 0.81 (LG13.822/LG10.709) to 0.14 (*LG13.826*/*LG14.2172*), consistent with the hypothesis that the genes were inserted into the *S*-locus region at different times. This finding supports the idea that the *S*-locus in *S. henryi* originated through a stepwise duplication process, as found in several other distylous species (e.g. Potente et al., 2022; Huu et al., 2020; Henning et al. 2023; Yang et al. 2023).

Comparative analysis and syntenic plotting between the two sub-genomes (haplotypes) indicated that *S*. *henryi* likely originated from auto-tetraploidization as illustrated in Fig. 7. The S-morph theoretically produces three types of diploid gametes, *SS, Ss* and *ss*, whereas the L-morph produces only *ss* gametes. When *SS* or *Ss* gametes fuse with *ss* gametes from the L-morph, the offspring are S-morph. And when *ss* gametes fuse with *ss* gametes, the offspring are L-morph. It has been suggested that *SS* genotypes are not viable in distylous *Primula* and other heterostylous groups (Kurian and Richards, 1997; Richards 1997). However, this is clearly not the case as viable *SS* plants are reported in both *Primula* (Yuan et al. 2019) and *Turnera* (Shore and Barrett 1985; Shore et al. 2019). Although *S. henryi* can theoretically produce both *Ss* and SS gametes, self- and intra-morph incompatibility (Deng, 2007) should prevent the formation of *SSSS* genotypes, although our genomic data indicates that *SS* gametes are likely viable since an *ssSS* individual was genotyped in our study.

**Figure 7.**
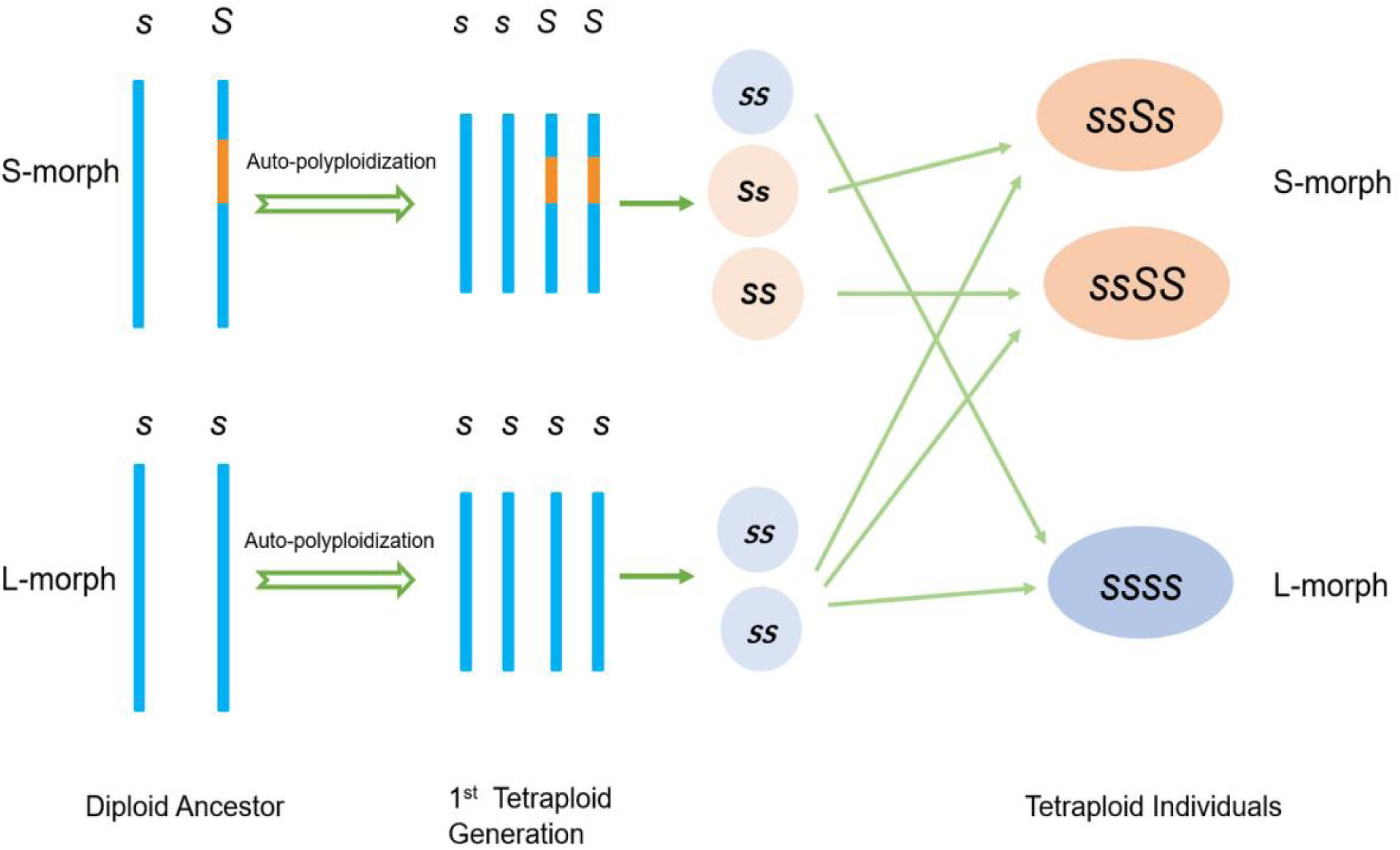
Postulated process for the formation of the tetraploid *Schizomussaenda henryi* from its diploid ancestor and genotypes of gametes and tetraploid individuals of the long- and short-styled morphs.

Our analysis of *Ks* values revealed that the most recent WGD event in *S*. *henryi* occurred at ca. 3.6 Mya and was represented by a peak at *Ks*=0.05596, which overlapped with the divergence peak for sub-genomes A and B (*Ks*=0.05606) (Fig. 3), further supporting that the genome of *S. henryi* had originated from auto-tetraploidization. This timing (3.6 Mya) was later than the divergence of *S. henryi* and *M. pubescens*, which was estimated to be ca. 7.15 Mya according to the *Ks* peak, and ca. 8 Mya based on an earlier phylogenetic analysis (Duan et al., 2018).

Estimates of *Ks* values between *S*-locus genes and their paralogs in *Primula* revealed that they had asynchronous origins, and that the chronological sequence was consistent with results inferred from molecular phylogenetic analysis (Potente et al., 2022). In *S*. *henryi*, the four *S*-locus genes exhibited *Ks* values ranging from 0.14 to 0.81, which corresponds to duplication ages of ca. 9.1 to 52.6 Myr. This finding suggests that the formation of the *S*-locus by stepwise duplication significantly predates the polyploidization event that occurred in *S. henryi,* which is positioned as the basal clade of the tribe Mussaendeae. Molecular phylogenetic evidence indicates that Mussaendeae split from other tribes of Rubiaceae ca. 47.3 Mya during the middle Eocene (95% HPD = 26.6–66.3 Mya) (Duan et al., 2018), and an earlier study by Bremer & Eriksson (2009) estimated the divergence time of Mussaendeae from other tribes to be 54.9 Mya. These estimates imply that the origin of the *S*-locus supergene controlling distyly may have begun as early as the divergence of Mussaendeae from their immediate ancestors and has been maintained at both diploid and tetraploid levels as a functioning outbreeding mechanism.

## Materials and Methods

### Study species, sample preparation and measurements

*Schizomussaenda henryi* (Hutch.) X. F. Deng & D. X. Zhang, the sole species in the genus, is an erect shrub or small tree distributed in southern China, northern Vietnam, Thailand, and Burma (Deng and Zhang, 2008). The golden flowers and white petaloid calyx-lobe make the species of significant horticultural value. The species is distylous and possesses self- and intra-morph incompatibility (Deng, 2007). We collected fresh leaves for genome sequencing from one L-morph individual and one S-morph individual growing in the South China Botanical Garden, Guangzhou, China which had been transplanted from a natural population at Mengla, Yunnan province, southwest China. Leaves were frozen in liquid nitrogen immediately and preserved for genome DNA extraction.

We collected floral buds and these were dissected in the lab, and style length and anther position measured using digital calipers (573-S, Mitutoyo, Japan) following the methods of Shore et al. (2019) and Luo et al. (2023), to facilitate sampling for RNA-sequencing. We divided flower development into early- and late-stages based on the elongation process of styles, corolla tubes and the relative position of stigma and anthers. Fresh styles and androecia from the two stages were collected in each morph: (1) Early-stage, ca. 5-8 mm corolla tube length, with stigmas and anthers of similar height; (2) Late-stage, ca. 14-18 mm corolla-tube length; with separation between stigma and anther (Fig. S1A). We collected styles by cutting them off at the top of the ovary, and the adnate filaments were collected with the corolla tube (anthers and corolla lobes removed). We sampled four biological replicates for each stage per morph with each replicate from a different individual from the population at Mengla, Yunnan Province. Thus, 32 samples (16 from the L-morph and 16 from the S-morph) were preserved in liquid nitrogen for transcriptomic sequencing.

### Karyotype analysis and genome size estimation

For chromosome counting, fresh shoot tips were collected from a single L-morph and S-morph plant and were the same individuals used for genome sequencing. After treatment in paradichlorobenzene saturated solution for 3 h, shoot tips were fixed in Carnoy’s Fluid (absolute ethanol: glacial acetic acid = 3: 1) for 30 min, washed by 95 % ethanol and pure water, and then treated in 1 mol/L HCl to dissociate the cells. After being rinsed in distilled water for three times, we stained the samples in Fuchsin solution for 2 h. The shoot tips were then crushed on a glass slide, which was covered and observed under an Olympus BX-41 microscope.

For genome size estimation, we chopped fresh leaves in pre-chilled OTTO buffer (containing RNase) using a sharp razor blade to release nuclei. Flow cytometry analysis followed procedures outlined by Dolezel and Bartos (2005), with *Zea mays* (genome size 2.1 Gb) chosen as the internal standards. We used GenomeScope v2.0 (Ranallo-Benavidez et al., 2020) and Jellyfish v2.2 (Marçais and Kingsford, 2011) for *k*-mer analysis and heterozygosity estimation, with parameters *k*=21, p=4.

### PacBio and Hi-C sequencing

We extracted genomic DNA from leaves using the DNeasy Plant Mini Kit (Qiagen, Germany) according to the manufacturer’s instructions. We assessed DNA quality and concentration by Qubit 2.0 Fluorometer (Life Technologies, USA) before sequencing. Then AMPure PB beads (PacBio 100-265-900, PacBio, USA) were employed for DNA purification to obtain high-quality genome DNA for library construction. We prepared size-selected SMRTbell libraries using SMRTbell® Express Template Prep Kit 2.0 (PacBio 101-853-100, PacBio, USA) and sequenced using the PacBio Sequel II system (PacBio, USA) by Biomarker Tech. (Beijing China).

We fixed leaves in 1% formaldehyde for HiC library construction, followed by adding glycine to quench the crosslinking reaction. Leaf tissues were then grounded in liquid nitrogen, and HindIII restriction endonuclease used to digest the cross-linked DNA. DNA ends were labeled with biotin-14-dCTP, and blunt-end ligation of the cross-linked fragments performed to form chimeric junctions. DNA fragments tagged with biotin were enriched with streptavidin magnetic beads and Hi-C libraries amplified through PCR and sequenced on an Illumina Novaseq 6000 platform with 2×150 bp reads.

### Transcriptome sequencing and expression analyses

Following the manufacturer’s instructions, we extracted total RNA from frozen samples with the aid of HiPure Universal RNA Mini Kit (Magen, Guangzhou, China). Before sequencing, we assessed RNA quality by Agilent Bioanalyzer 2100 system (Agilent Technologies, CA, USA). RNA samples with RIN (RNA Integrity Number) > 7.0 and OD 260/280 > 2.0 were quantified for PacBio library preparation, Illumina library construction, and sequencing.

We used the TruSeq RNA Sample Prep Kit (IlluminaRS-122-2002, Illumina, USA) for poly(A) mRNA isolation, first-strand and second-strand cDNA synthesis, fragment and adapter ligation, as well as cDNA library preparation. We sequenced the libraries from different tissues on an Illumina Novaseq 6000 platform using 150bp pair-end sequencing. We obtained clean reads by removing adaptors, undetermined bases (poly-N), and lowquality reads (those with more than 50% bases with a Q-value ≤ 20).

We then mapped clean reads to the genome of *S. henryi* using HISAT2 (Kim et al., 2019) and quantified expression abundances of unigenes with the aid of featureCounts (Liao et al., 2014). We calculated FPKM (fragments per kilobase of exon per million mapped fragments) using StringTie (Shumate et al., 2022) to represent the relative expression levels of each transcript. A gene was considered as ‘expressed’ if the FPKM in at least one tissue was above 1.0. We performed differential expression analysis by the DESeq2 package (ver. 1.26.0) in R and statistical test results (*p*-values) were corrected using the Benjamini-Hochberg false discovery rate (FDR). We assigned genes with FDR ≤ 0.05 and FC (fold change) ≥ 2 (log 2 FC ≥ 1 or log 2 FC ≤ -1) between two transcriptomes as DEGs.

### Genome assembly and chromosome anchoring

We assembled the genome of *S. henryi* with HiFiasm (version 0.16.1) using PacBio HiFi reads (CCS mode), with HiC data incorporated (Cheng et al., 2021). The quality of the initial assembly was assessed by BUSCO (v.5.1, with database primates_odb, 13,780 genes), based on expected gene content from near-universal single-copy orthologs (Manni et al., 2021).

We performed quality control for Hi-C data using *fastp* to remove the adaptors, trim reads and eliminate the low-quality bases (Chen, 2023). Clean Hi-C reads pairs were mapped to the *S. henryi* genome using BWA (0.7.17) with default parameters. Paired reads with mates mapped to a different contig were used in the Hi-C associated scaffolding. LACHESIS (Burton et al., 2013) was further applied to perform contig grouping, intra-group orienting and sorting, and we used *Coffea canephora* as a reference for chromosome assignment. We visualized the assemblies and manually corrected with the aid of Juicebox (Durand et al., 2016) and used YaHS (Zhou et al., 2023) to generate the final genome assembly after the edit was completed. We conducted chromosome anchoring using Biomarker Tech. (Beijing China).

### Repetitive sequence prediction

Based on the chromosome-level assemblies, we integrated *de novo* and homology-based approaches to identify repeat sequences in the genome of *S. henryi*. We used RepeatModeler v2.0.1 (Flynn et al., 2020) to construct the *de novo* repeat library, and the EDTA (Extensive de novo TE Annotator, Ou et al. 2019, 2022) pipeline to predict transposable elements (TEs). TEsorter (v.1.4.6) (Zhang et al., 2022) was used to refine the classification of TEs. Then we used RepeatMasker v4.1.0 (Smit, 2013) to identify, annotate and softmask the repetitive elements with the combined database of the Repbase, *de novo* repeat library, and the TE library.

To analyze TE enrichment, we calculated TE densities using 5kb windows calculated for the *S*-locus and flanking regions in the S-morph genome with the aid of TBtools-II (Chen et al., 2023).

### Gene prediction and annotation

We performed gene structure prediction incorporating *de novo*, homology-based, and transcriptomic data-based annotation workflows. We used Augustus (http://augustus.gobics.de), SNAP (Korf, 2004) and GENEID (Blanco et al., 2007) for *de novo* prediction. Homology-based prediction was based on GEMOMA (Keilwagen et al., 2018), using the genomes from *Coffea canephora*, *Chiococca alba, Arabidopsis thaliana*, and *Oryza sativa* as references. We employed TRINITY (Haas et al., 2013) to assemble the RNA-seq data and the results were provided to PASA for gene structure modeling.

We combined results from different workflows to generate the prediction gene set with the aid of EVM (Haas et al., 2008). Genes with TE insertions were identified and removed by TEsorter (v.1.4.6), followed by filtering with AGAT (v.1.2.0) (Dainat, 2024) to eliminate incomplete genes. We evaluated the accuracy and completeness of gene predictions by BUSCO based on the “protein” model (Manni et al., 2021).

We performed functional annotation for protein-coding genes using different databases, including Swissprot (https://web.expasy.org/docs/swiss-prot_guideline.html), PFAM (http://xfam.org/), eggNOG (http://eggnogdb.embl.de/), GO (http://geneontology.org/page/go-database), and KEGG (http://www.genome.jp/kegg/) through the eggNOG-mapper (v.2) pipeline (Cantalapiedra et al., 2021). We further filtered the *S*-locus genes by their annotations, and those which did not have a hit in the Swissprot or could not be assigned to any gene families were excluded.

### Sub-genome reconstruction and analyses

Based on the chromosome-level assemblies, we conducted collinearity analysis using JCVI, a python version of MCscan (Tang et al., 2008), and divided the chromosomes into homoeologous pairs, with each group containing 11 chromosomes. We employed SubPhaser (v1.2.5) (Jia et al., 2022) to identify sub-genome-specific k-mers, but this approach failed to divide the genome into two groups with equal chromosome numbers (see Results). Single-copy genes were identified with the aid of OrthoFinder (v2.5.4) (Emms and Kelly, 2019) in each sub-genome and in the closely related *Mussaenda pubescens* (Luo et al., unpublished data). We constructed a phylogenetic tree and the genetic distance for each chromosome was calculated through the OrthoFinder pipeline, with parameters -M msa -T iqtree. We assigned the 11 chromosomes with closer distances to *M. pubescens* to sub-genome A and another 11 chromosomes were assigned to sub-genome B (Luo et al., 2024). We performed the genomic collinearity analysis between the two sub-genomes by JCVI utility libraries and genomic syntenic depth for these species pairs was calculated through the ’jcvi.compara.synteny depth’ pipeline in JCVI with default parameters.

To detect potential sub-genome dominance, sub-genome A and B were compared. We calculated the number of genes, mRNA, exons, introns and CDS of each sub-genome using TBtools-II and compared the contents of repetitive sequences between the two sub-genomes. We focused our BUSCO analysis on missing genes in the sub-genomes using a highly conserved gene database (2,326 genes). Sub-genome-specific missing BUSCO genes were analyzed through functional enrichment using GSEA (Gene Set Enrichment Analysis) software (v. 4.2.3) (Subramanian et al., 2005).

We identified homoeologous gene pairs in the tetraploid genome of *S. henryi* using OrthoFinder (version 2.4.0) (Emms and Kelly, 2019) and selected the high-quality genome of *Coffea canephora* as the diploid reference. We performed the all-against-all blastp search with an E-value cutoff of 1e-5. We used orthologous genes showing 1:1:1 relationship in *C. canephora* and the two sub-genomes of *S. henryi* to calculate the *Ka* and *Ks* values.

The compared the expression of syntenic gene pairs (homoeologous genes) to detect homoeolog expression dominance (Xu et al., 2019). Genes pairs with Fold Change greater than two were defined as dominant gene pairs with genes expressed higher in dominant gene pairs considered dominant genes.

### Whole-genome duplication analyses

To detect and compare the WGD events in *S. henryi* and three other species [*M. pubescens* (Luo et al. 2025) and *C. canephora* (Rubiaceae), and *Thymus quinquecostatus* (Lamiaceae)], we used all-against-all BLASTP to perform one-to-one comparisons of homologs (e-value cut-off 1*10^-6^). We chose a sample from Lamiaceae as it is a sister clade of Gentianales, the order that includes Rubiaceae. Previous studies found that *C. canephora* and *T. quinquecostatus* had experienced one and two rounds of WGD, respectively (Denoeud et al., 2014; Sun et al., 2022). The comparison was also performed between sub-genomes A and B of *S. henryi,* with the methods described above. We calculated *Ks* values (synonymous substitution rates) of the collinear orthologous gene pairs with WGDI (Whole-Genome Duplication Integrated analysis) using the ’Ks’ pipeline (Sun et al., 2021). We generated A *Ks* dotplot of gene anchor pairs through the ’block’ pipeline, and the ’KsPeaks’ pipeline was employed for the distribution analysis of the *Ks* median value. Prominent *Ks* peaks indicate potential whole-genome duplication events.

### Identification and analysis of distyly-related genes

To identify candidate genes involved in the regulation of style-stamen dimorphism, we compared gene expression between the L- and S-morphs at each developmental stage for styles and filaments (corolla tubes). We focused first on genes that were exclusively expressed in the S-morph (FPKM<1.0 in L-morph). Other DEGs that could be potentially related to style or filament elongation, such as those involved in BR, GA and/or Auxin metabolic pathways, were also selected for analysis.

We mapped PacBio sequencing data of the S-morph to the assembled genome with the aid of BWA (Li, 2013). The .sam file was converted to .bam format and sorted by SAMtools (Li et al., 2009), and sequencing depth was calculated by PanDepth (https://github.com/HuiyangYu/PanDepth/). Assuming hemizygosity of the S-locus region, the sequencing depth for the *S*-locus would be expected to be about half of that for flanking regions which are not hemizygous.

We visualized the genome sequences and gene annotations and these were compared in IGV (Integrative Genomics Viewer, v.2.4.10) to examine if candidate genes were S-morph specific, i.e. gene sequences only present in the S-morph but absent from corresponding regions in the L-morph. We performed whole genome resequencing to verify the putative *S*-locus regions based on read-coverage analysis. We collected samples of young leaves from the natural population at Mengla, Yunnan Province and three individuals of each morph were subsequently sequenced on the Illumina Novaseq 6000 platform using 150bp pair-end sequencing, with an expected coverage of 10X. We aligned clean data to our genome assemblies using BWA, and the resulting .bam file was sorted and indexed by SAMtools and mapping coverage computed by *samtools coverage*. We visualized and compared aligned sequences with the aid of IGV.

We selected *S*-locus genes that were reported from earlier studies of distylous *Primula, Gelsemium, Turnera, Linum*, and *Fagopyrum* (cited in Introduction) to search against L- and S-morph genomes using the “Find Best Homology” module of TBtools-II (Chen et al., 2023), which integrates reciprocal BLAST and phylogenetic analyses. Genes with the closest relationship to the input genes were selected as putative paralogs and their expression data compared between the two morphs to evaluate their potential role as candidate distyly genes.

### Origin and evolution of *S*-locus genes

We identified paralogs of *S*-locus genes and flanking genes in L- and S-morph genomes using the “Find Best Homology” module of TBtools-II. We calculated *Ks* values (rate of synonymous substitution) between each *S*-locus gene and its paralogous gene using the “Simple Ka/Ks Calculator” module in TBtools-II to estimate molecular divergence between alleles. To characterized divergence of regions flanking the *S*-locus, we extracted and aligned sequences 20kb upstream and 20kb downstream of the *S*-locus in the S-morph genome, and corresponding regions in the L-morph genome, using MEGA11 (Tamura et al., 2021). We then cut into windows of 200bp with our own *perl* scripts and calculated Evolutionary Distances (EDs) of each window using the K2P (Kimura two-parameter) method implemented in MEGA11, with a transitions + transversions substitution model.

## Acknowledgements

We are grateful to Professor Elena Conti (University of Zurich) for providing a preprint of work on homostylous polyploid *Primula*, Guanshen Liu from Biomarker Tech. (Beijing) for technical support, and Hongyan Li for lab assistance.

## Author contributions

DX.Z., S.C.H.B., ZL.L. and MM.S. designed and coordinated the study. TY.T., MM.S., DX.Z. and ZL.L. provided the funding. SS.J., BQ.X. and SR.G. participated in sample collection, preparation and lab works. ZL.L., MM.S, ZT. Z, and Y.Z. led the bioinformatics analyses. TT.D assisted in phylogenetic analyses. LB.J. contributed to the genome analyses. L.G. and XF.D. helped in data processing. ZL.L. wrote the manuscript. S.C.H.B., DX.Z. and ZL.L. revised the manuscript. All authors discussed the results and approved the manuscript.

## Data availability

All data supporting the results of this study are included in the manuscript and its supplemental files. Genome assemblies and raw sequence data were deposited in the CNSA database (https://db.cngb.org), and annotation files have been uploaded to Figshare (https://figshare.com), which will be available to public upon acceptance of the manuscript.

## Funding

This work was supported by the National Natural Science Foundation of China (31970252, 32170232, 32070222, 32271613), the Guangdong Provincial Special Fund for Natural Resource Affairs on Ecology and Forestry Construction (GDZZDC20228704), and the Initial Funding for Doctoral Research from Shandong University of Technology (Grant No. 4041/422047).

---

**Table S1.**
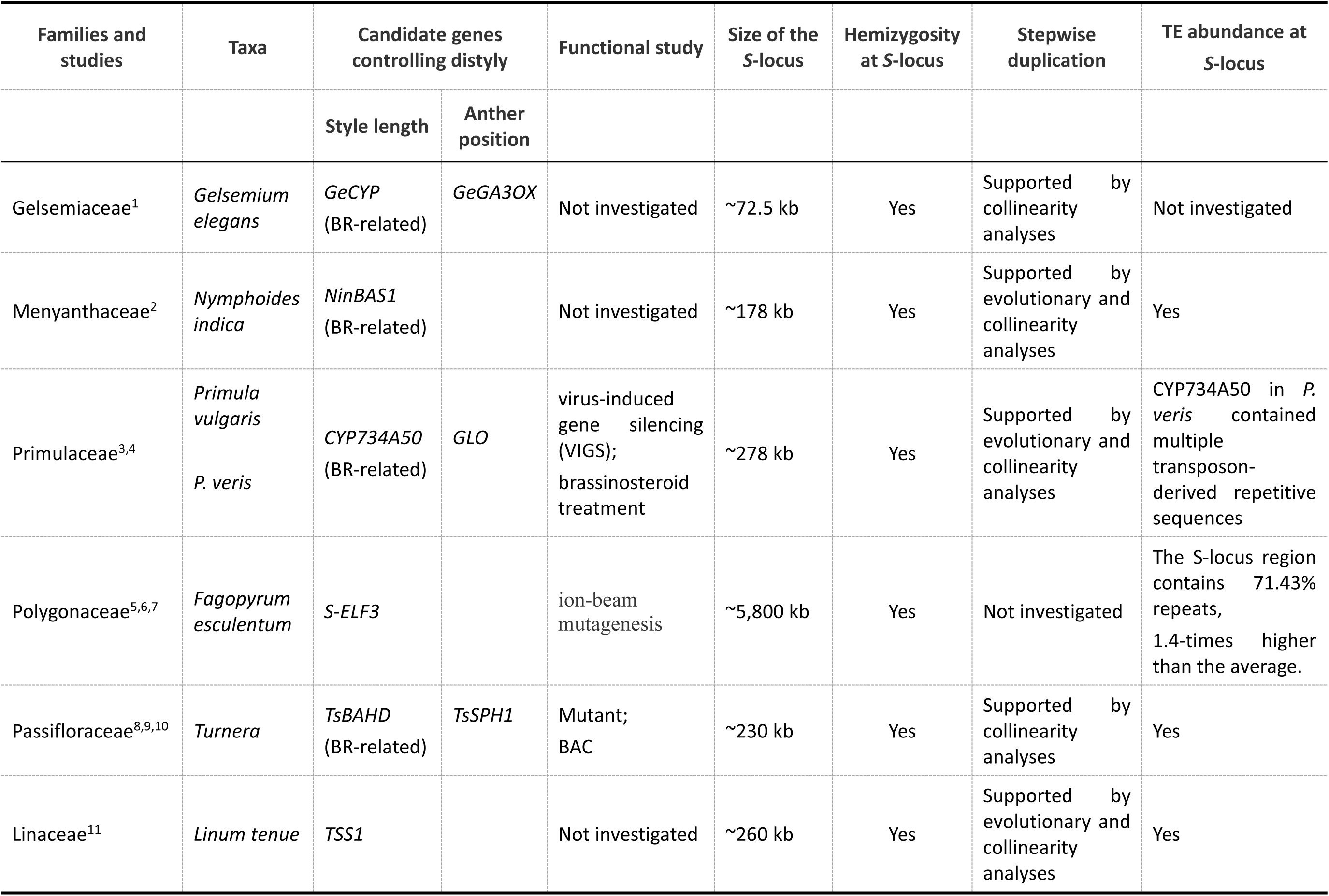
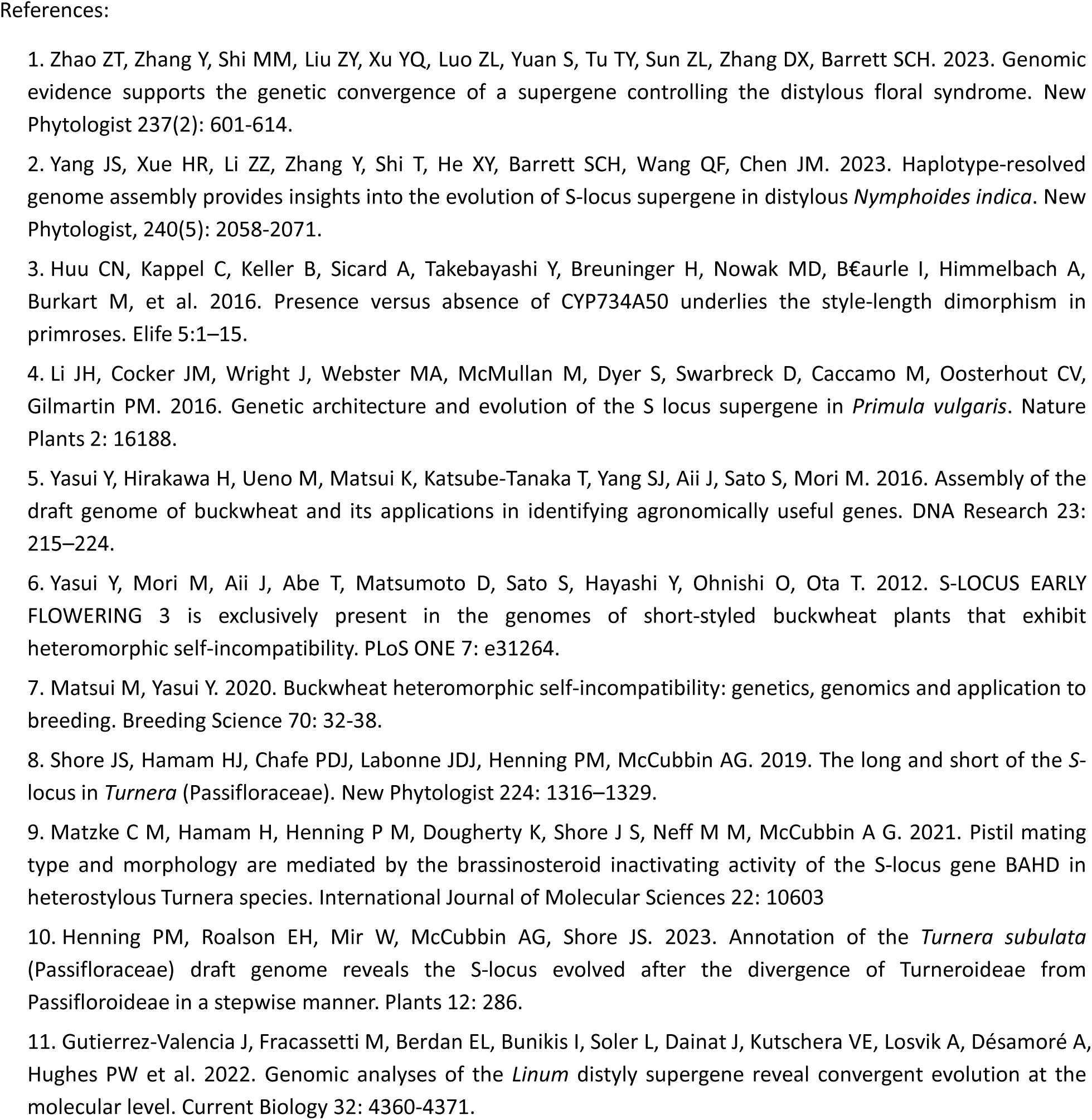
*S*-locus information from published studies on distylous species in six angiosperm families.

| Families and studies | Taxa | Candidate genes controlling distyly |  | Functional study | Size of the S-locus | Hemizyosity at S-locus | Stepwise duplication | TE abundance at S-locus |
| --- | --- | --- | --- | --- | --- | --- | --- | --- |
|  |  | Style length | Anther position |  |  |  |  |  |
| Gelsemiaceae<sup>1</sup> | *Gelsemium elegans* | *GeCYP* (BR-related) | *GeGA3OX* | Not investigated | ~72.5 kb | Yes | Supported by collinearity analyses | Not investigated |
| Menyanthaceae<sup>2</sup> | *Nymphoides indica* | *NinBAS1* (BR-related) |  | Not investigated | ~178 kb | Yes | Supported by evolutionary and collinearity analyses | Yes |
| Primulaceae<sup>3,4</sup> | *Primula vulgaris*<br>*P. veris* | *CYP734A50* (BR-related) | *GLO* | virus-induced gene silencing (VIGS); brassinosteroid treatment | ~278 kb | Yes | Supported by evolutionary and collinearity analyses | *CYP734A50* in *P. veris* contained multiple transposon-derived repetitive sequences |
| Polygonaceae<sup>5,6,7</sup> | *Fagopyrum esculentum* | *S-ELF3* |  | ion-beam mutagenesis | ~5,800 kb | Yes | Not investigated | The S-locus region contains 71.43% repeats, 1.4-times higher than the average. |
| Passifloraceae<sup>8,9,10</sup> | *Turnera* | *TsBAHD* (BR-related) | *TsSPH1* | Mutant; BAC | ~230 kb | Yes | Supported by collinearity analyses | Yes |
| Linaceae<sup>11</sup> | *Linum tenue* | *TSS1* |  | Not investigated | ~260 kb | Yes | Supported by evolutionary and collinearity analyses | Yes |

**Table S2.** BUSCO assessment for Hi-C assembly of the genomes of the long- and short-styled morphs of *Schizomussaenda henryi*.

| BUSCO parameters | Percentage (%) |  |
| --- | --- | --- |
|  | L-morph | S-morph |
| Complete BUSCOs | 98.6 | 98.8 |
| Complete and single-copy BUSCOs | 10.6 | 10.4 |
| Complete and duplicated BUSCOs | 88.0 | 88.4 |
| Fragmented BUSCOs | 0.5 | 0.3 |
| Missing BUSCOs | 0.9 | 0.9 |

**Table S3.** Results of functional annotations of genes for the long- and short-styled morphs of

| Database | L-morph |  | S-morph |  |
| --- | --- | --- | --- | --- |
|  | No. of genes | Percentage of genes | No. of genes | Percentage of genes |
| KEGG | 17,558 | 33.77% | 17,573 | 33.41% |
| GO | 34,190 | 65.75% | 34,500 | 65.59% |
| Uniprot | 24,134 | 46.41% | 24,526 | 33.41% |
| Annotated | 44,880 | 86.32% | 45,327 | 86.17% |
| Unannotated | 7,075 | 13.61% | 7,273 | 13.83% |
| Total | 51,955 | - | 52,600 | - |

**Table S4.**
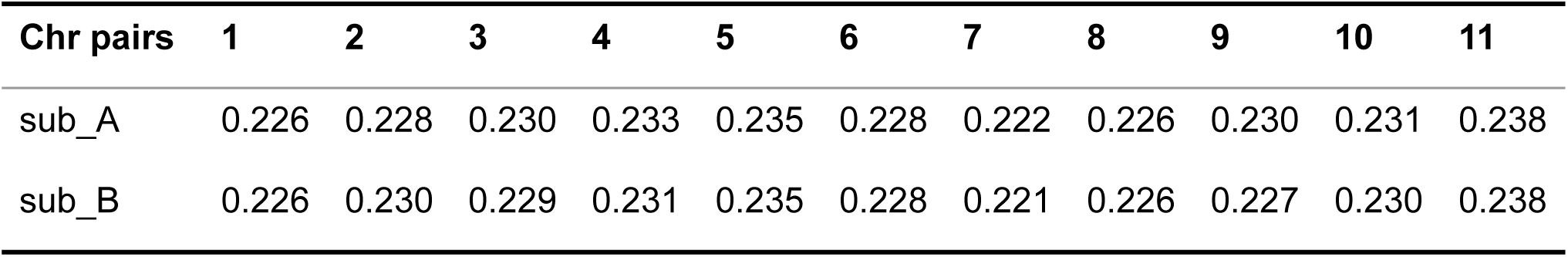
Mean *Ka/Ks* ratios of protein-coding genes in each chromosome of sub-genomes A and B in the S-morph of *Schizomussaenda henryi*.

**Figure S1.**
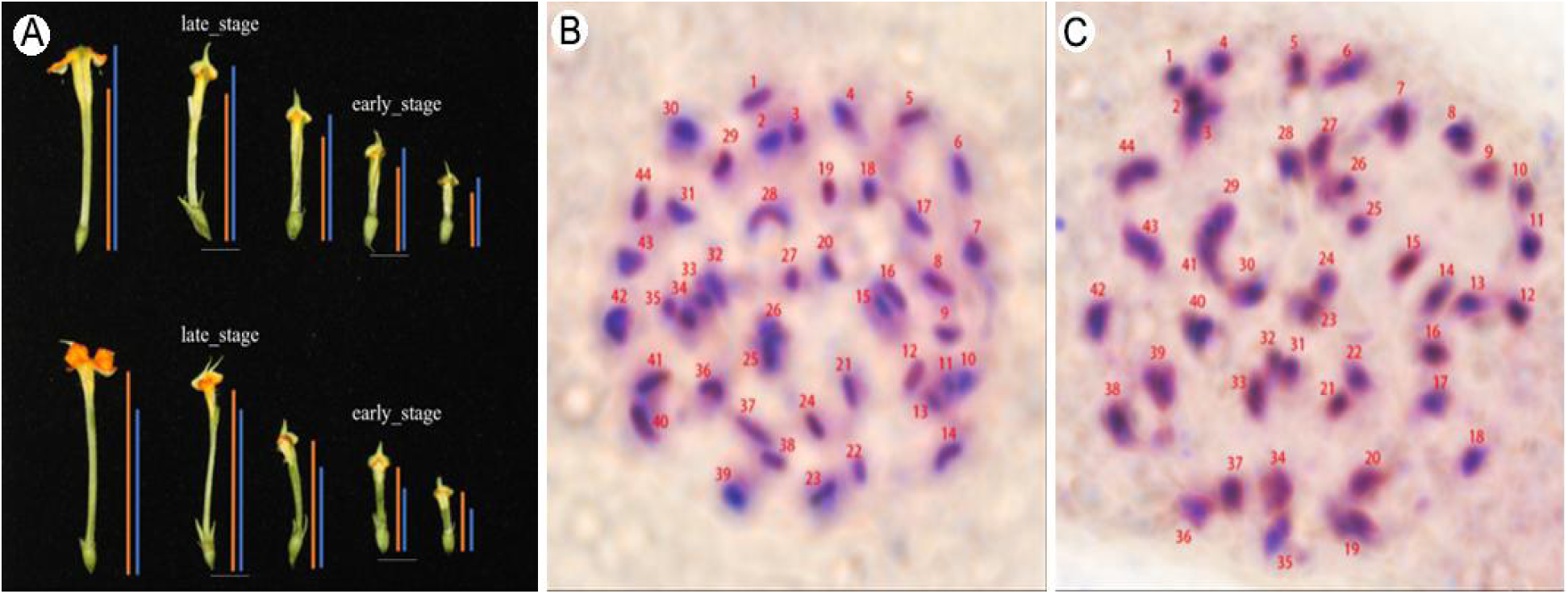
A. Dissection of flowers of the long- and short-styled morphs of *Schizomussaenda henryi*, illustrating the two developmental stages for RNA_seq sampling; B, C. Microscopic examination of stem tip cell chromosomes of L-morph (B) and S-morph (C) *S. henryi.* Chromosomes are numbered 1-44.

**Figure S2.**
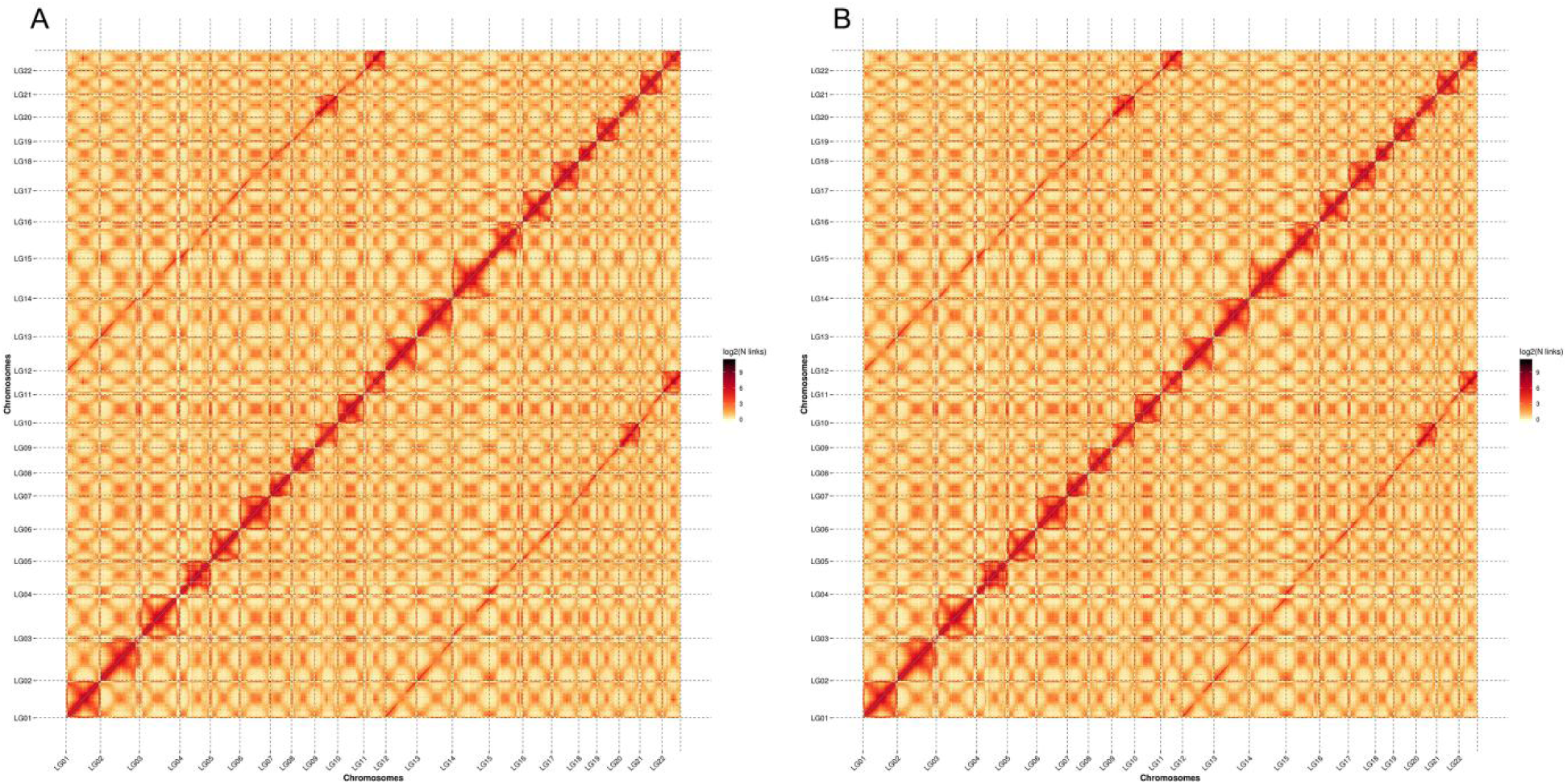
Heatmaps of Hi-C interaction density for the L-morph (A) and S-morph (B) genomes of *Schizomussaenda henryi*. The interaction density for each pair of windows (resolution: 300 kb) is measured by the number of supporting Hi-C reads.

**Figure S3.**
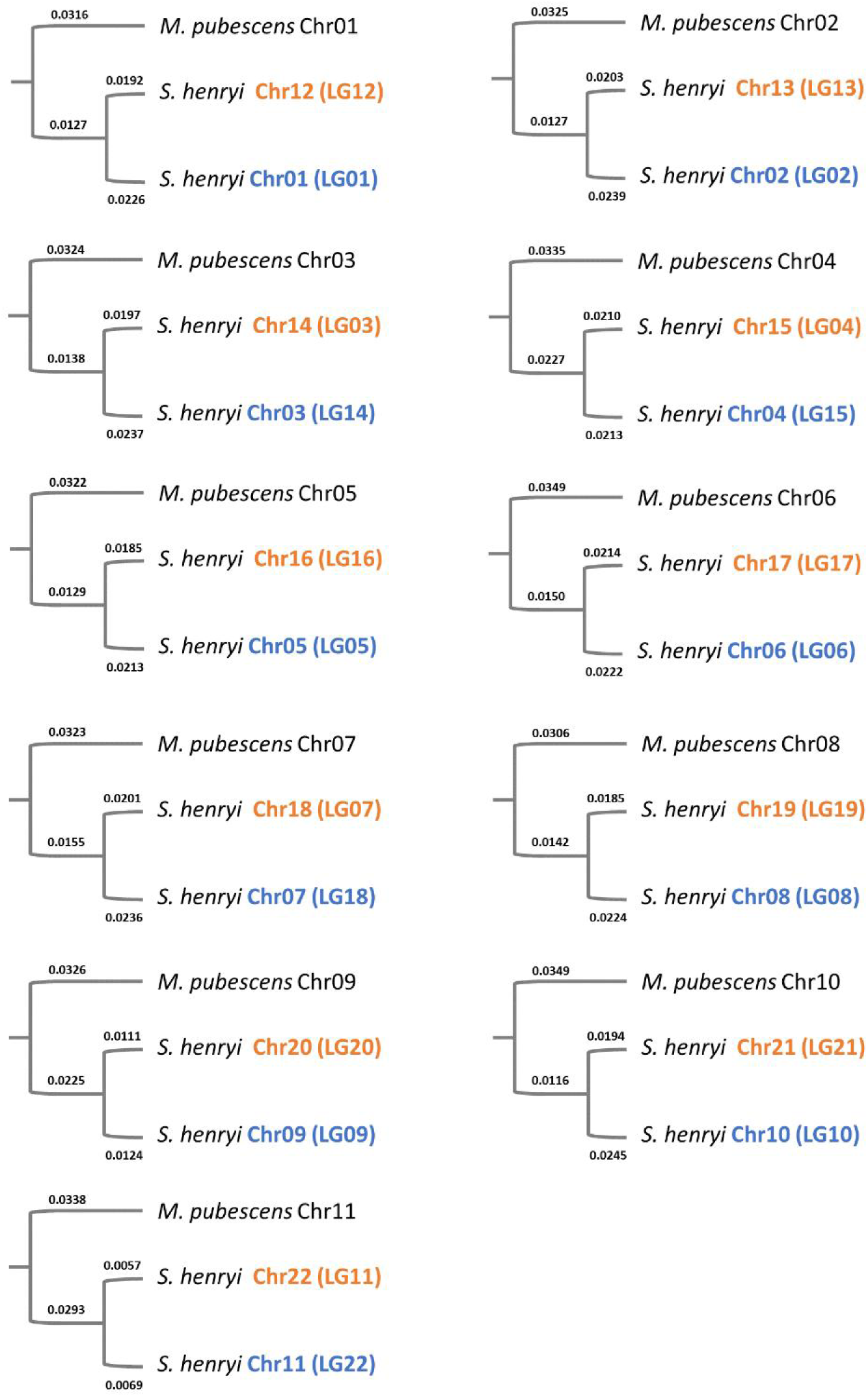
Measures of genetic distance between *Schizomussaenda henryi* and the close relative *Mussaenda pubescens* support the assignment of chromosomes to the A (blue) and B (orange) sub-genomes.

**Figure S4.**
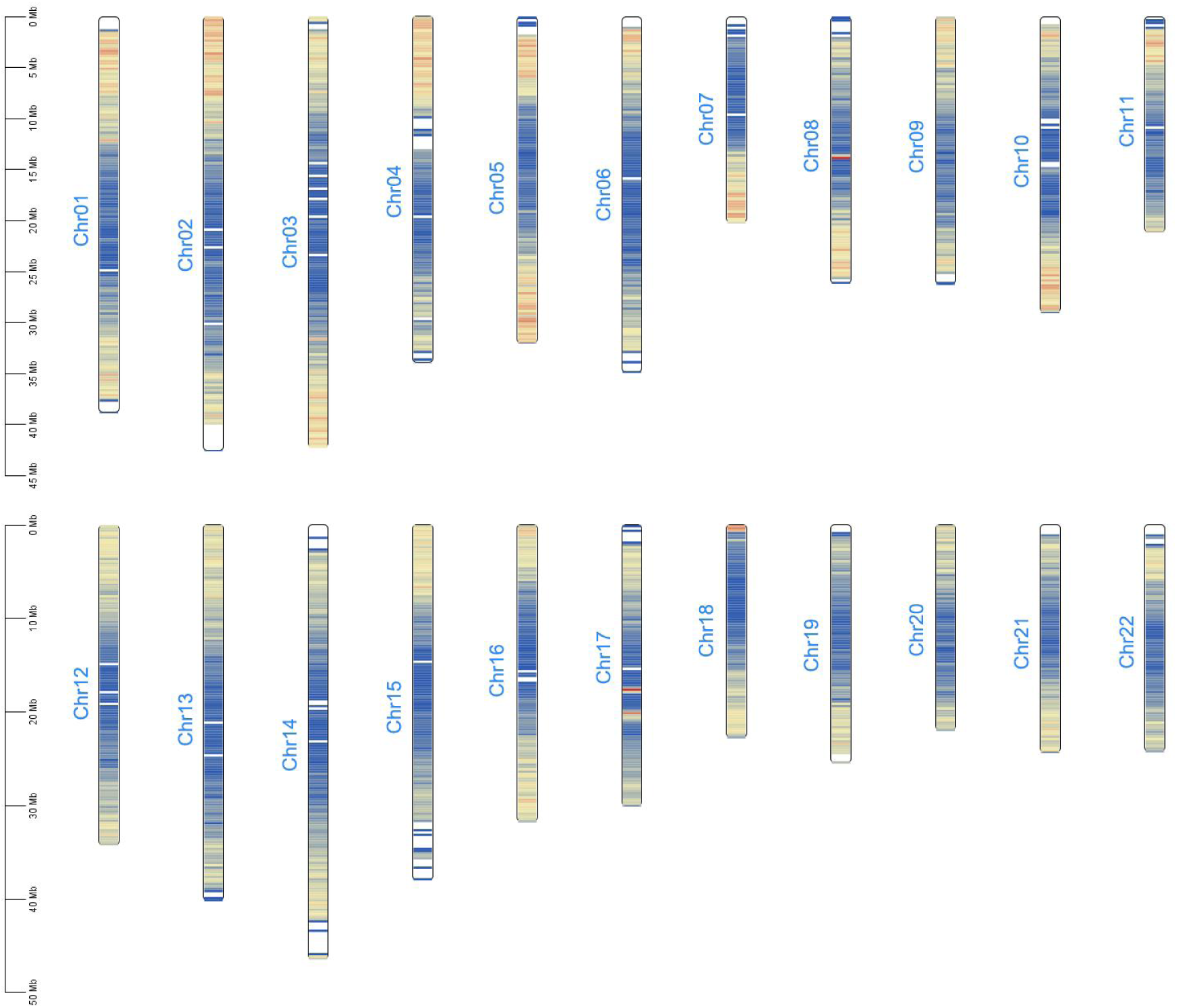
Gene density distributions on each chromosome in the genome of the short-styled morph of *Schizomussaenda henryi*.

**Figure S5.**
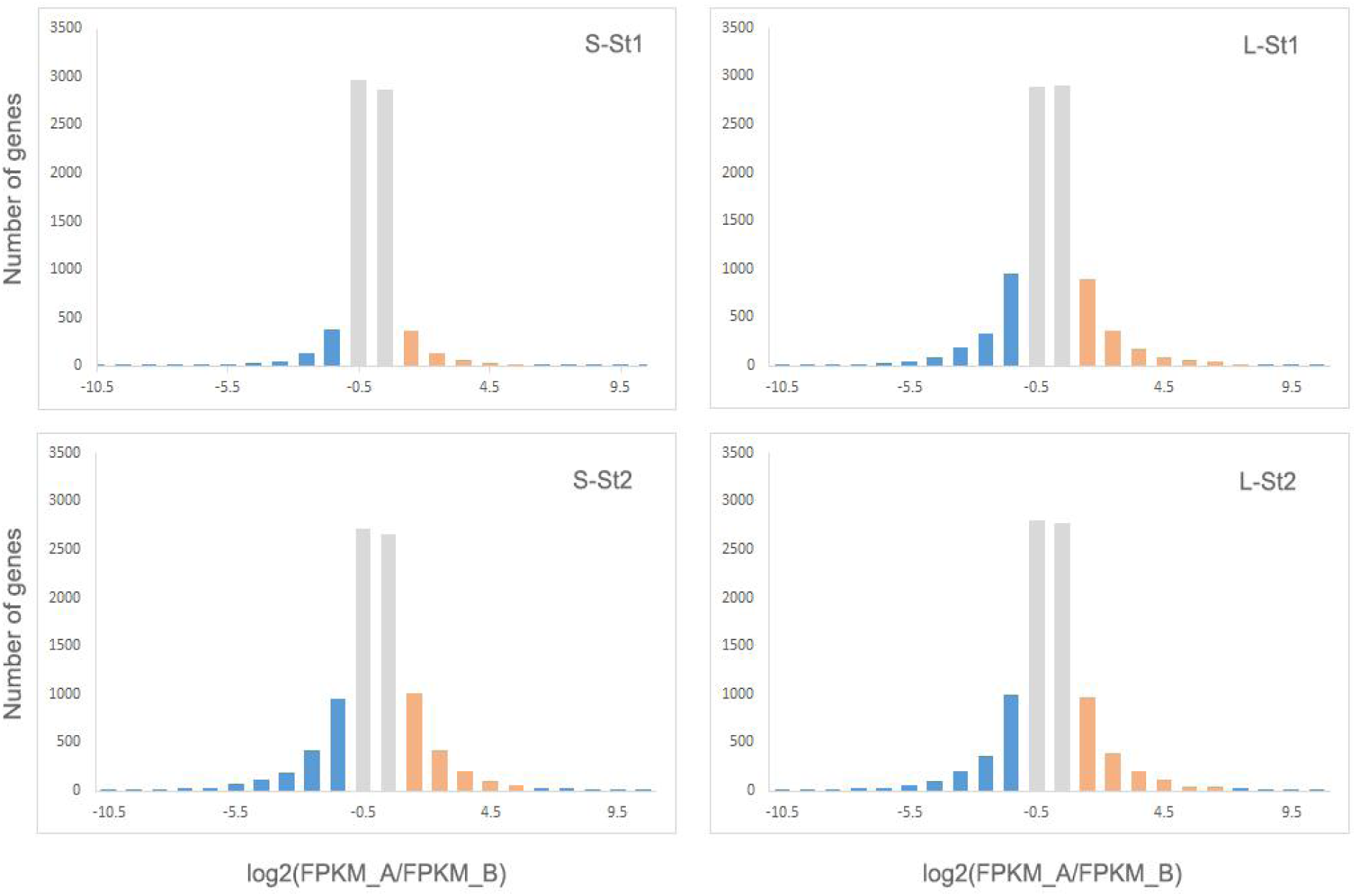
Histograms showing the expression of homoeologous genes in style tissue of the short-styled -morph of *Schizomussaenda henryi* at different developmental stages. Log2(FPKM_A/FPKM_B) indicates the degree of expression difference of homoeologous gene pairs. St1, early stage; St2, late stage.

**Figure S6.**
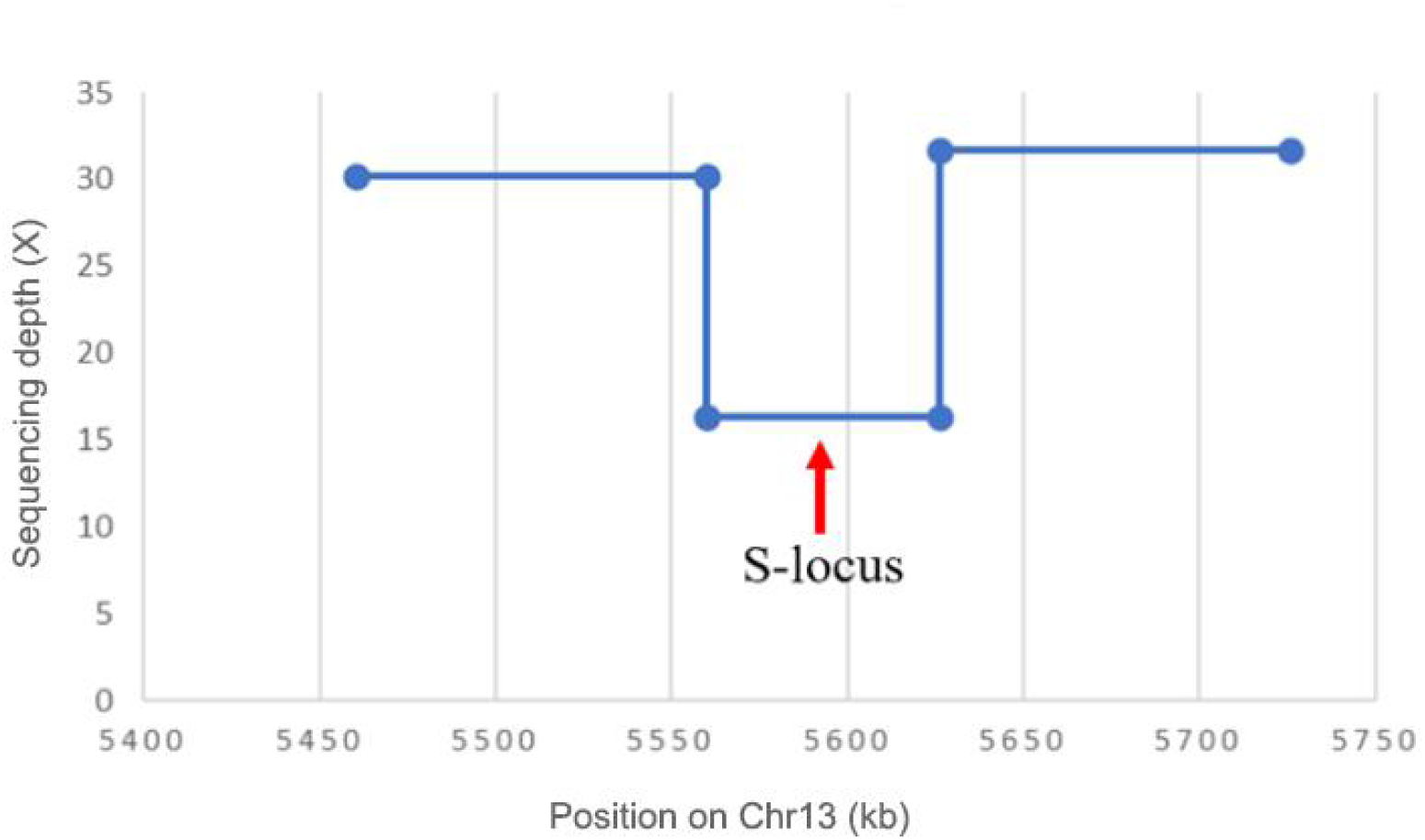
The sequencing depth of the *S*-locus in this study for the S-morph genome of *Schizomussaenda henryi*.

**Figure S7.**
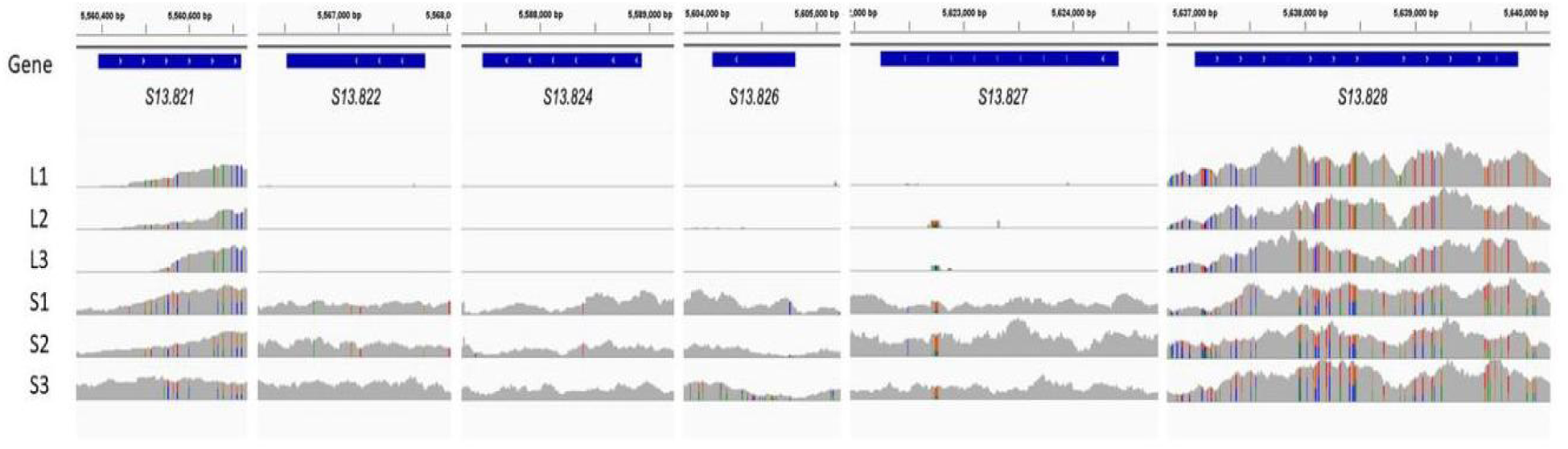
Alignments of resequencing data of long- and short-styled morphs at the *S*-locus of *Schizomussaenda henryi* (visualized by IGV). L1-L3 represent the three L-morph individuals, and S1-S3 the three S-morph individuals.

**Figure S8.**
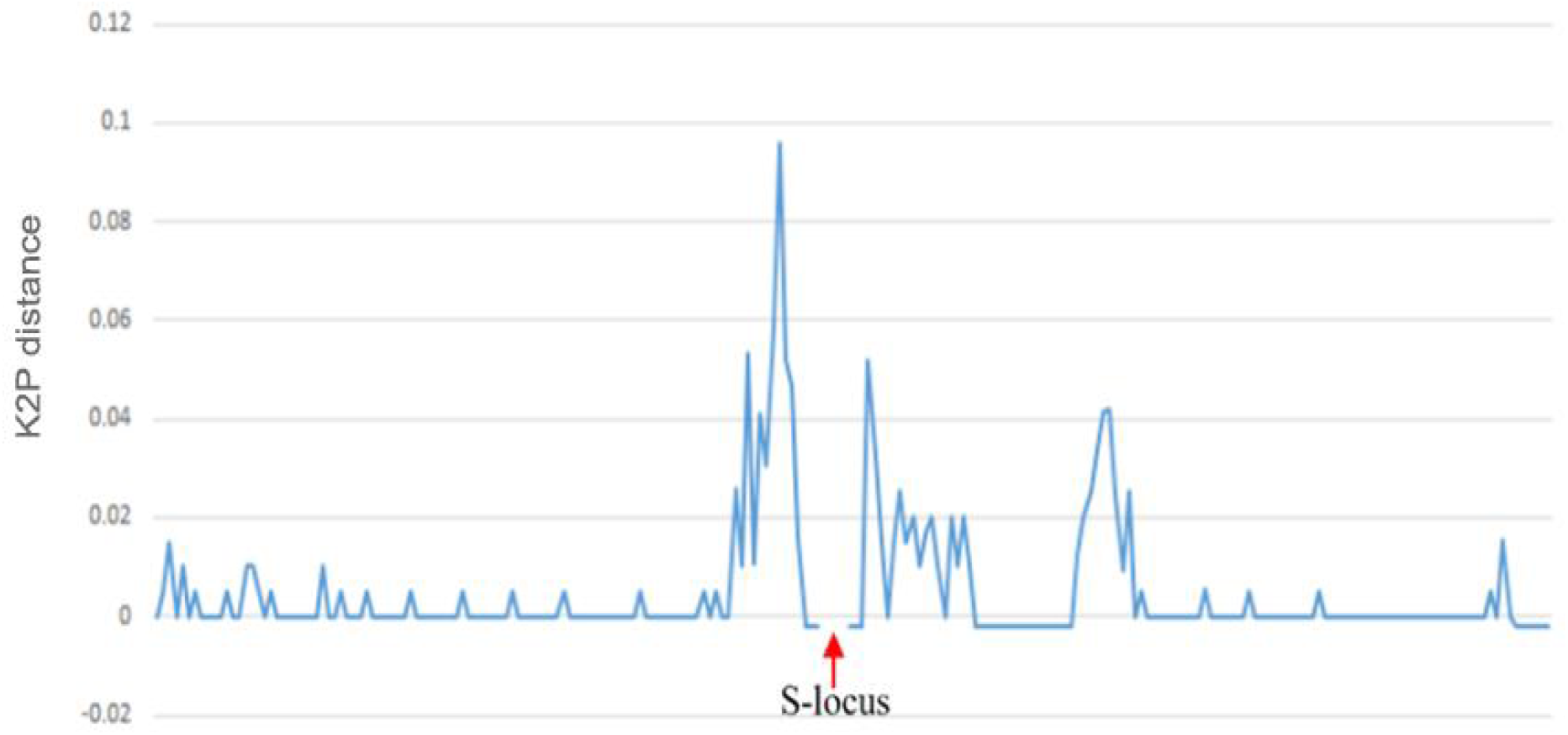
Evolutionary divergence (Kimura two-parameter distance – K2P distance) of sequences flanking the *S*-locus in the short-styled morph of *Schizomussaenda henryi*.

**Figure S9.**
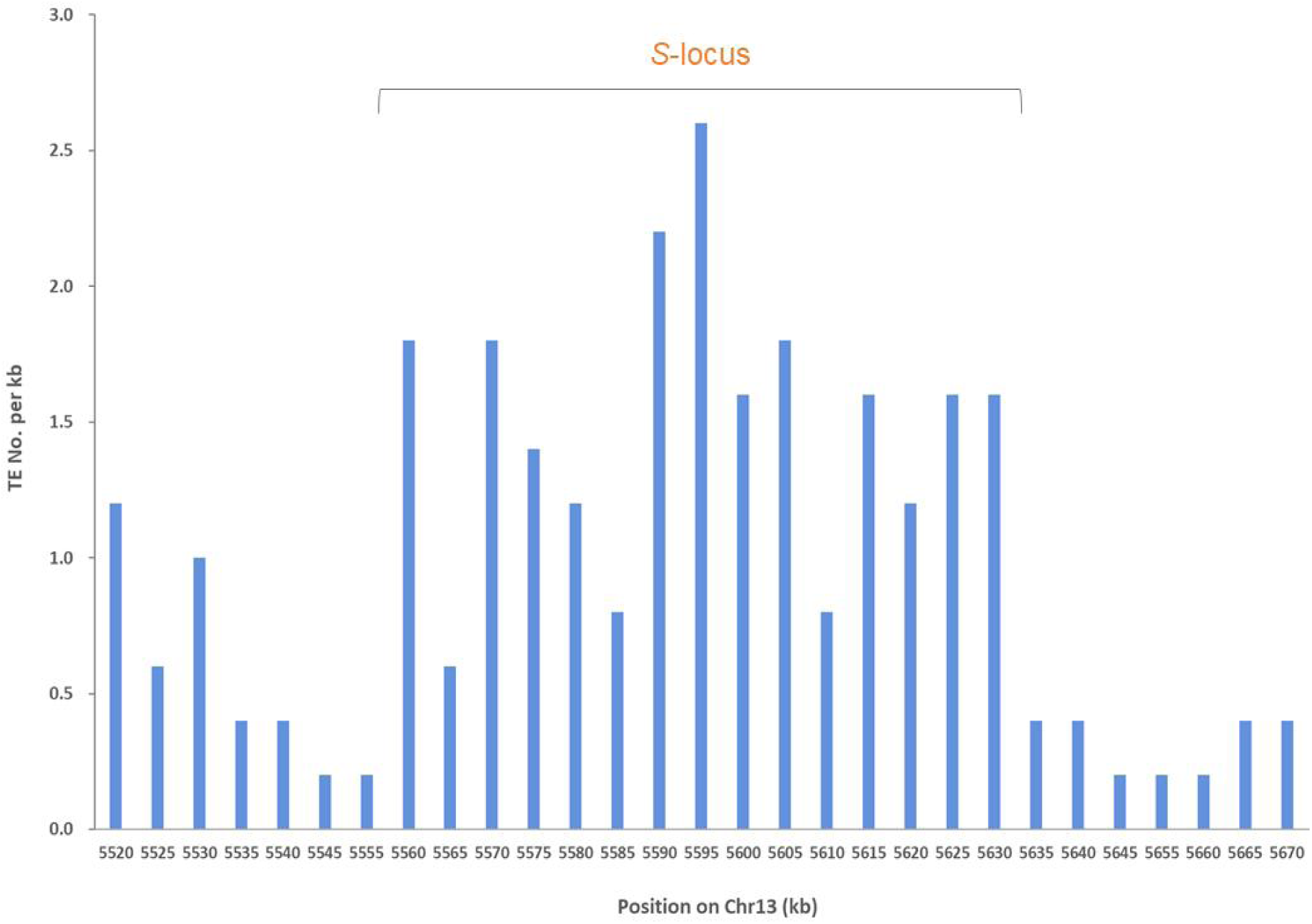
TE density in the *S*-locus region and flanking regions (5 kb for each window) of *Schizomussaenda henryi* (5,562 - 5,637 kb).

## Notes

### Competing Interest Statement

The authors have declared no competing interest.

