## Supplementary Data for "Genetic architecture of the *S*-locus supergene revealed in a tetraploid distylous species"

Supplementary Data 1

SubPhaser results for S-morph genome

| Group | subgenome | bootstrap |
|---|---|---|
| LG01 | SG1 | 100 |
| LG12 | SG1 | 100 |
| LG02 | SG1 | 100 |
| LG13 | SG1 | 100 |
| LG03 | SG1 | 100 |
| LG14 | SG1 | 100 |
| LG15 | SG1 | 100 |
| LG05 | SG1 | 100 |
| LG16 | SG1 | 100 |
| LG06 | SG1 | 100 |
| LG17 | SG1 | 100 |
| LG07 | SG1 | 100 |
| LG18 | SG1 | 100 |
| LG19 | SG1 | 100 |
| LG09 | SG1 | 100 |
| LG10 | SG1 | 100 |
| LG21 | SG1 | 100 |
| LG11 | SG1 | 100 |
| LG22 | SG1 | 100 |
| LG04 | SG2 | 100 |
| LG08 | SG2 | 100 |
| LG20 | SG2 | 93 |

SubPhaser results for L-morph genome

| Group | subgenome | bootstrap |
|---|---|---|
| LG01 | SG1 | 100 |
| LG12 | SG1 | 99 |
| LG02 | SG1 | 99 |
| LG13 | SG1 | 99 |
| LG03 | SG1 | 99 |
| LG14 | SG1 | 99 |
| LG04 | SG1 | 99 |
| LG05 | SG1 | 100 |
| LG16 | SG1 | 99 |
| LG17 | SG1 | 98 |
| LG07 | SG1 | 99 |
| LG18 | SG1 | 99 |
| LG19 | SG1 | 100 |
| LG09 | SG1 | 99 |
| LG10 | SG1 | 99 |
| LG21 | SG1 | 96 |
| LG11 | SG1 | 99 |
| LG22 | SG1 | 97 |
| LG15 | SG2 | 77 |
| LG06 | SG2 | 87 |
| LG08 | SG2 | 99 |
| LG20 | SG2 | 97 |

Subgenome division based on phylogeny

| Group | sub_A | Group | sub_B |
|---|---|---|---|
| LG01 | Chr01 | LG12 | Chr12 |
| LG02 | Chr02 | LG13 | Chr13 |
| LG14 | Chr03 | LG03 | Chr14 |
| LG15 | Chr04 | LG04 | Chr15 |
| LG05 | Chr05 | LG16 | Chr16 |
| LG06 | Chr06 | LG17 | Chr17 |
| LG18 | Chr07 | LG07 | Chr18 |
| LG08 | Chr08 | LG19 | Chr19 |
| LG09 | Chr09 | LG20 | Chr20 |
| LG10 | Chr10 | LG21 | Chr21 |
| LG22 | Chr11 | LG11 | Chr22 |

Supplementary Data 2: Subgenome missing BUSCOs

| BUSCOs missing in subA | BUSCOs missing in subB |
|---|---|
| 101717at71240 | 101557at71240 |
| 107927at71240 | 101717at71240 |
| 110565at71240 | 105398at71240 |
| 115938at71240 | 110565at71240 |
| 116194at71240 | 110801at71240 |
| 116773at71240 | 113616at71240 |
| 117781at71240 | 116194at71240 |
| 117864at71240 | 117864at71240 |
| 119695at71240 | 118793at71240 |
| 124041at71240 | 122159at71240 |
| 124055at71240 | 124055at71240 |
| 12447at71240 | 129938at71240 |
| 126919at71240 | 134860at71240 |
| 127152at71240 | 138740at71240 |
| 128778at71240 | 14548at71240 |
| 129938at71240 | 147003at71240 |
| 133348at71240 | 147683at71240 |
| 133861at71240 | 14824at71240 |
| 134342at71240 | 148977at71240 |
| 134860at71240 | 149408at71240 |
| 135371at71240 | 14960at71240 |
| 135744at71240 | 150253at71240 |
| 138237at71240 | 151593at71240 |
| 139195at71240 | 152308at71240 |
| 140271at71240 | 152354at71240 |
| 140511at71240 | 15546at71240 |
| 14071at71240 | 15721at71240 |
| 142995at71240 | 172984at71240 |
| 14634at71240 | 177383at71240 |
| 147683at71240 | 18835at71240 |
| 147895at71240 | 23252at71240 |
| 14878at71240 | 24187at71240 |
| 148950at71240 | 27136at71240 |
| 148977at71240 | 27601at71240 |
| 149408at71240 | 28528at71240 |
| 152308at71240 | 29774at71240 |
| 15422at71240 | 29956at71240 |
| 156253at71240 | 312at71240 |
| 164300at71240 | 32704at71240 |
| 16664at71240 | 32894at71240 |
| 167878at71240 | 33509at71240 |
| 172525at71240 | 35592at71240 |
| 17405at71240 | 35686at71240 |
| 18383at71240 | 39655at71240 |
| 18672at71240 | 40951at71240 |
| 20162at71240 | 4609at71240 |
| 22198at71240 | 48937at71240 |
| 23252at71240 | 50420at71240 |
| 2400at71240 | 50574at71240 |
| 24187at71240 | 5406at71240 |
| 24315at71240 | 56188at71240 |
| 25215at71240 | 57567at71240 |
| 28072at71240 | 59001at71240 |

| BUSCOs missing in subA | BUSCOs missing in subB |
|---|---|
| 28090at71240 | 59224at71240 |
| 29774at71240 | 60492at71240 |
| 29956at71240 | 62825at71240 |
| 31173at71240 | 63750at71240 |
| 32894at71240 | 65474at71240 |
| 34577at71240 | 65567at71240 |
| 35585at71240 | 66769at71240 |
| 37280at71240 | 67815at71240 |
| 40995at71240 | 67911at71240 |
| 41392at71240 | 71319at71240 |
| 41521at71240 | 72570at71240 |
| 42883at71240 | 72635at71240 |
| 43180at71240 | 74501at71240 |
| 48585at71240 | 76106at71240 |
| 4933at71240 | 769at71240 |
| 55281at71240 | 78542at71240 |
| 59224at71240 | 81777at71240 |
| 60160at71240 | 83990at71240 |
| 60277at71240 | 86443at71240 |
| 60659at71240 | 91118at71240 |
| 6543at71240 | 91264at71240 |
| 65567at71240 | 91524at71240 |
| 68243at71240 | 96666at71240 |
| 68663at71240 | 9742at71240 |
| 68795at71240 | 98387at71240 |
| 70034at71240 | 98964at71240 |
| 707at71240 | |
| 71626at71240 | |
| 72570at71240 | |
| 74085at71240 | |
| 76961at71240 | |
| 82519at71240 | |
| 83990at71240 | |
| 85405at71240 | |
| 85704at71240 | |
| 86443at71240 | |
| 89090at71240 | |
| 91118at71240 | |
| 93922at71240 | |
| 94224at71240 | |
| 95431at71240 | |
| 96736at71240 | |
| 98964at71240 | |

Supplementary Data 3: BLAST results using the S-locus gene sequences against the four haplotype-resolved assemblies

| | | Haplotype 1 | | |
|---|---|---|---|---|
| query id | refer id | identity (%) | e-value | bit score |
| S13.822 | ctg0331 | 92.97 | 0 | 1866 |
| S13.824 | ctg0179 | 88.77 | 0 | 808 |
| S13.826 | ctg0294 | 92.29 | 0 | 1085 |
| S13.827 | ctg0218 | 90.33 | 3.51E-106 | 390 |

| | | Haplotype 2 | | |
|---|---|---|---|---|
| query id | refer id | identity (%) | e-value | bit score |
| S13.822 | ctg0052 | 93.15 | 0 | 1858 |
| S13.824 | ctg0109 | 88.77 | 0 | 808 |
| S13.826 | ctg0331 | 92.29 | 0 | 1085 |
| S13.827 | ctg0244 | 90.33 | **3.45E-106** | 390 |

| | | Haplotype 3 | | |
|---|---|---|---|---|
| query id | refer id | identity (%) | e-vaue | bit score |
| S13.822 | ctg0046 | 100.00 | 0 | 2366 |
| **S13.824** | **ctg0046** | **100.00** | **0** | **2687** |
| S13.826 | ctg0046 | 100.00 | 0 | 1410 |
| **S13.827** | **ctg0046** | **100.00** | **0** | **4021** |

| | | Haplotype 4 | | |
|---|---|---|---|---|
| query id | refer id | identity (%) | e-vaue | bit score |
| S13.822 | ctg0045 | 100.00 | 0 | 2366 |
| **S13.824** | **ctg0045** | **100.00** | **0** | **2687** |
| S13.826 | ctg0045 | 100.00 | 0 | 1410 |
| **S13.827** | **ctg0045** | **100.00** | **0** | **4021** |
